# Peptide-mediated inhibition of the transcriptional regulator Elongin BC induces apoptosis in cancer cells

**DOI:** 10.1101/2022.11.04.515028

**Authors:** Sabrina Fischer, Van Tuan Trinh, Clara Simon, Lisa Marie Weber, Ignasi Formé, Andrea Nist, Gert Bange, Frank Abendroth, Thorsten Stiewe, Wieland Steinchen, Robert Liefke, Olalla Vázquez

**Affiliations:** Institute of Molecular Biology and Tumor Research (IMT), University of Marburg, 35043, Germany; Department of Chemistry, University of Marburg, 35043 Marburg, Germany; Protein Analysis Unit, Biomedical Center (BMC), Faculty of Medicine, Ludwig-Maximilians-University (LMU) Munich, 82152 Martinsried, Germany; Genomics Core Facility, Institute of Molecular Oncology, Member of the German Center for Lung Research (DZL), University of Marburg, 35043 Marburg, Germany; Center for Synthetic Microbiology (SYNMIKRO), University of Marburg, 35043 Marburg, Germany; Department of Hematology, Oncology, and Immunology, University Hospital Giessen and Marburg, 35043 Marburg, Germany

## Abstract

Inhibition of protein-protein interactions (PPIs) via designed peptides is an effective strategy to interfere with their biological functions. The Elongin BC heterodimer (ELOB/C) is involved in transcription elongation and protein turnover by PPIs that involve the so-called BC-box. ELOB and ELOC are commonly upregulated in cancer and essential for cancer cell growth, making them attractive drug targets. However, no strategy has been established to inhibit their functions in cells, so far. Here, we report a peptide that mimics a high-affinity BC-box and tightly binds to the ELOB/C dimer (*k*_D_ = 0.45 ± 0.03 nM). Our peptide blocks the association of ELOB/C with its interaction partners, both in vitro and in the cellular environment. Cancer cells treated with this peptide inhibitor show decreased cell viability, altered cell cycle and increased apoptosis. Therefore, our work proposes that blocking the BC-box binding pocket of ELOB/C is a feasible strategy to impair the function of the ELOB/C heterodimer and inhibit cancer cell growth. Our peptide inhibitor promises novel mechanistic insights into the biological function of the ELOB/C dimer and offers a starting point for therapeutics linked to ELOB/C dysfunction.

## Main

Protein-protein interactions (PPIs) play a central role in life^1^. They enable spatial proximity among proteins, which results in the formation of multiprotein complexes. These quaternary structures dominate biological processes such as translation, transcription, signal transduction, and many more. The highly complex cellular network of PPIs is commonly known as the “interactome”. Aberrant alterations of the interactome often cause diseases, including cancer^2^. Consequently, many efforts have been undertaken to influence PPIs^2^. In contrast to enzymes, whose substrate-binding pockets typically can easily be targeted, PPIs have long been considered undruggable due to their difficult topology. However, recent progress has demonstrated that also PPIs can be disrupted efficiently^2^. Both small molecules and peptides have been reported to successfully interfere with PPIs^3^. Along these lines, the current “peptide tidal wave” placed peptides as promising chemical biology tools for function interference^4^. Indeed, peptides represent a sweet spot between small molecules and biologics, which is particularly suitable for directly targeting PPIs. Peptides can simulate crucial secondary structures along the PPI interface, achieving efficient disruption. Thus, peptides overcome the issues derived from the typical poor surface architecture of PPIs. In addition, their straightforward synthesis enables a broad structural diversity, which entails selectivity, versatility and high potency without compromising biocompatibility. Therefore, peptides are valuable tools for proof-of-concept studies to elucidate novel therapeutic targets. The recent large increase in structural information, based on experimental^5^ and computational methods^6^, opened up an entirely new space for drug discovery based on PPI modulation.

A classical example of PPIs is the Elongin BC heterodimer (ELOB/C). ELOB/C is composed of two proteins, namely the 118 amino acid protein Elongin B (ELOB) and the 112 amino acid protein Elongin C (ELOC). ELOB/C forms a hydrophobic surface that allows the interaction with a short alpha-helical motif, called BC-box. The sequence of the BC-box is commonly described as P/T/S-L-X-X-X-C/S/A-X-X-X-ϕ_7_, but is found in various variants^8^. The only amino acid that is irreplaceable is an N-terminal leucine, which reaches deep into the binding groove of ELOB/C^9^. ELOB/C interacts with a large variety of partners, including SOCS (suppressors of cytokine signalling) proteins, the Von-Hippel-Lindau tumour suppressor (VHL), the mediator subunit MED8, the transcriptional elongation factor ELOA, and many more^7^. Most ELOB/C interaction partners also possess a cullin box, allowing the formation of an E3 ubiquitin ligase complex. This complex consists of ELOB/C, cullin 2/5, and the BC-box containing proteins. These multiprotein associations are involved in the ubiquitination and, in turn, degradation of their target proteins^7^. Of special interest in this regard are the SOCS proteins, which play a major role in the immune system by regulating cytokine signalling pathways^10^. Besides its major role in regulating protein degradation, ELOB/C has also been implicated in protein stabilisation ^11^ and as a component of the transcription elongation factor SIII^12^. Together with the VHL, ELOB/C is an integral part of the cellular response to hypoxia^9,13^. ELOB/C has also been demonstrated to be associated with the PRC2-interacting protein EPOP^14-16^. This association has been implicated in the interplay between transcriptional repression by PRC2 and active transcription. Thus, the PPIs between the BC-box sequence and ELOB/C trigger the assembly of versatile multiprotein complexes that are involved in numerous cellular processes, both in the cytoplasm and in the nucleus.

Here, we report that ELOB/C is a putative drug target in cancer. Bioinformatic approaches based on public CRISPR screens suggested that cancer cells require ELOB/C for survival, which was further confirmed by knockdown experiments. To assess the suitability of ELOB/C as a drug target, we designed a peptide inhibitor that mimics the BC-box sequence of EPOP. Fluorescence polarization (FP) and hydrogen/deuterium exchange (HDX) mass spectrometry confirmed that our peptide tightly binds to the BC-box binding pocket of ELOB/C (*k*_D_ = 0.45 ± 0.03 nM). Our peptide can efficiently disrupt the interaction of ELOB/C with its interaction partners in vitro and cells, while its scrambled version as well as the mutated version do not display any effect. Viability assays together with clonogenicity experiments confirmed the superior effect of our peptide inhibitor compared to its variants in three different cancer cell lines. Importantly, we demonstrated that ELOB/C inhibition in living cells leads to perturbed gene expression of cancer-related pathways and increases apoptosis. Therefore, our work discovers the potential therapeutic role of ELOB/C and highlights its inhibition as a feasible strategy to impair cancer cell growth.

## Results

### CRISPR screens identify the ELOB/C dimer as a potential target for cancer therapy

To identify novel putative targets for cancer therapy, we investigated publicly available CRISPR screening data that were performed in multiple human cancer cell lines^17,18^. Typical oncogenes, such as MYC, have a strong negative average CRISPR score, while common tumour suppressors, such as PTEN, have a positive average CRISPR score (**Fig. 1a**). In this context, we also identified the genes for Elongin B (*ELOB*) and Elongin C (*ELOC*) to be required for the growth of most human cancer cell lines **(Fig. 1a)**. The roles of *ELOB* and *ELOC* are similar in cancer cell lines from all tissues, demonstrating the importance of these genes in most cancer cell types (**Fig. 1b**). Interestingly, although ELOB and ELOC form a heterodimer^12^, implying that they form a functional unit, the CRISPR scores of both proteins show hardly any correlation (**Extended Data Fig. 1A**) and the knockout of *ELOB* often has a stronger impact on cancer cells (**Fig. 1b**), leading to a more negative CRISPR score in most cancer cell types.

**Fig. 1:**
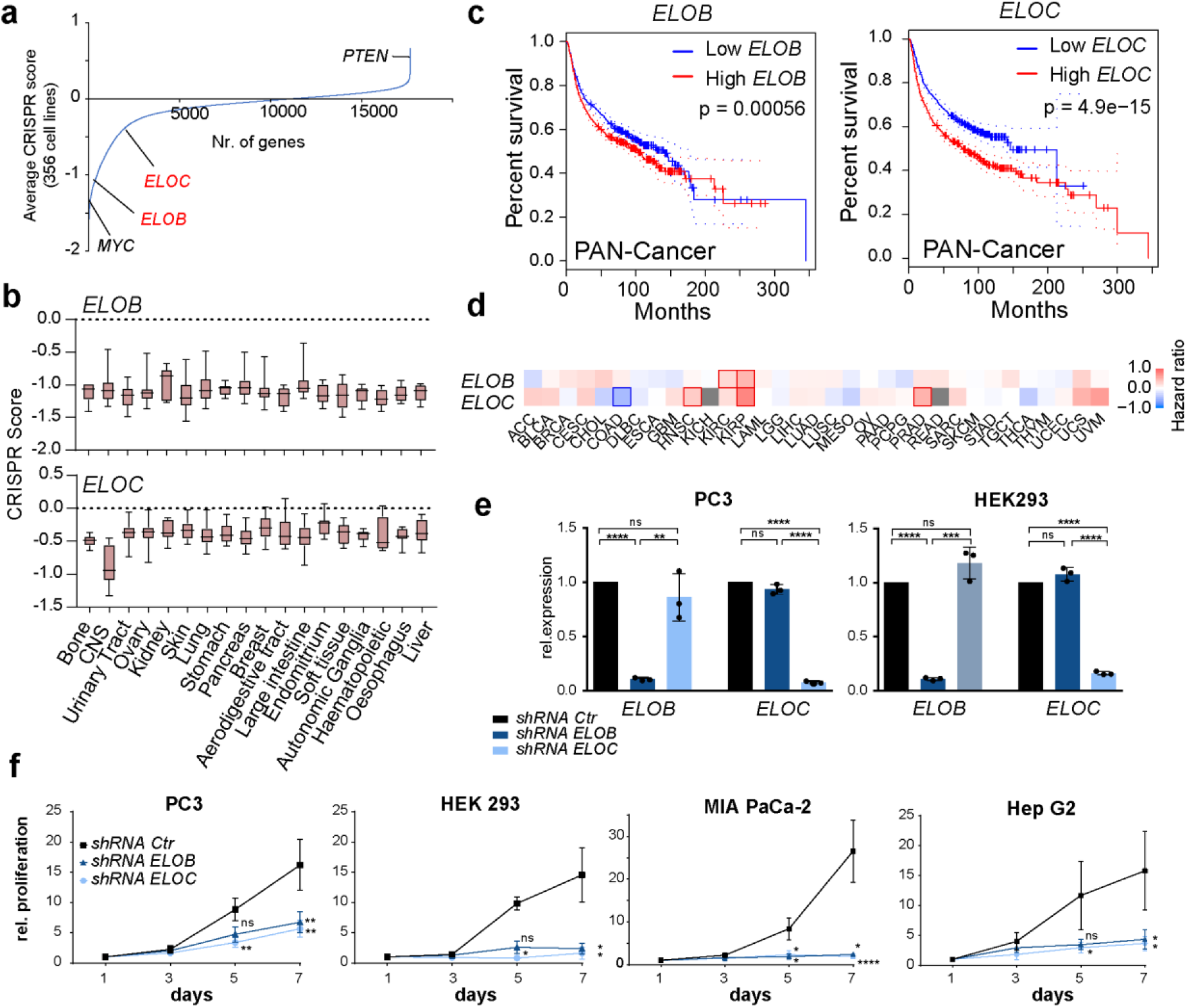
ELOB/C is essential for cancer cell growth. **a**, Genes sorted according to their average CRISPR score measured in 356 cancer cell lines. A negative score indicates low survival upon CRISPR knockout of the respective genes. **b**, Distribution of the CRISPR scores of *ELOB* and *ELOC* in cancer cell lines of various cancer types. The whisker-box plots represent the lower quartile, median and upper quartile of the data with 5% and 95% whiskers. **c**, Kaplan-Meier survival plots of disease-free survival based on expression from *ELOB* (left) and *ELOC* (right) in all cancer types. Data from TCGA^19^ and visualized by GEPIA^20^. **d**, Survival map of disease-free survival based on *ELOB* and *ELOC* expression in various cancer types, visualized by GEPIA^20^. **e**, RT-qPCR analysis of *ELOB* and *ELOC* gene expression after their knockdown in PC3 and HEK293 cells. Data are presented as mean ± s.d. For statistical analysis, an unpaired t-test with the assumption of Gaussian distribution and similar s.d. variance was conducted. Three independent replicates were used. **f**, Proliferation of cancer cells after knockdown of *ELOB* and *ELOC*. Data are presented as the mean ± s.d. At least three independent replicates were collected and a ratio paired t-test of paired samples was conducted on the 5^th^ and 7^th^ day. A Gaussian distribution was assumed as well as consistent ratios of paired values. n.s., *p* > 0.05; *, *p* < 0.05; **, *p* < 0.01; ***, *p* < 0.001; ****, *p* < 0.0001.

Investigation of gene expression data from TCGA (The Cancer Genome Atlas)^19,20^ demonstrates that *ELOB* and *ELOC* are both commonly upregulated in many cancer tissues (**Extended Data Fig. S1B**) and that high expression typically is associated with a worse prognosis, further supporting a relevant role of both proteins in cancer (**Fig. 1c**). However, looking at individual cancer types, the gene expression often correlates distinctly with the patient survival **(Fig. 1d**). Thus, these results suggest that although ELOB and ELOC are known to form a heterodimer, their roles in cancer appear to be nonoverlapping.

### Knockdown of Elongin B and C impairs essential cellular pathways in PC3 prostate cancer cells

The molecular reasons why ELOB and ELOC are essential for cancer cell growth are currently poorly understood. To gain insights into the role of the ELOB/C in cancer cells, we depleted *ELOB* and *ELOC* from HEK293 cells and five distinct cancer cell lines (PC3 prostate cancer cells, MIA PaCa-2 pancreatic carcinoma cells, HepG2 liver cancer cells, NCI-H23 lung cancer cells and SH-SY5Y neuroblastoma cells) using specific shRNAs. We confirmed the efficient knockdown of both *ELOB* and *ELOC* via RT-qPCR experiments in HEK293 and PC3 cells (**Fig. 1e**). Consistent with the results from the CRISPR screen, we found that knockdown of *ELOB* and *ELOC* led to a strongly reduced proliferation in all investigated cell lines (**Fig. 1f, Extended Data Fig. 1d**). Notably, ELOB and ELOC do not appear to be essential in every cell type. Previous work suggests that the deletion of *ELOB* in mouse ES cells had only minor consequences on the biological properties of the cells^14^.

Given that high *ELOB* and *ELOC* expression correlates with negative prognosis for prostate cancer patients (**Fig. 1d, Extended Data Fig. 1c**), we investigated the role of ELOB/C in further detail using the PC3 prostate cancer cell line as a model. We performed RNA-Seq experiments, after the depletion of *ELOB* and *ELOC* in these cells, in three replicates. Principal component analysis (PCA) of the obtained data demonstrated that the gene expression pattern after *ELOB* and *ELOC* knockdown cells strongly differed from that of the control cells, as expected (**Fig. 2a**). Knockdown of *ELOB* led to the dysregulation of 723 (*p* < 0.01, log2 fold change > 0.75) genes, while we observed 1,345 dysregulated genes upon knockdown of *ELOC* (**Fig. 2b**). Unexpectedly, the consequences of *ELOB* and *ELOC* knockdown were rather distinct, with only a small overlap of the differentially expressed genes (**Fig. 2c**). This finding suggests that ELOB and ELOC may have functions that are independent of each other. This hypothesis is further supported by the observation that ELOB and ELOC do not have identical cellular distributions, as assessed by immunofluorescence in HEK293 cells. Although both proteins are predominantly in the nucleus, ELOB also shows substantial staining in the cytoplasm (**Extended Data Fig. 2**).

**Fig. 2:**
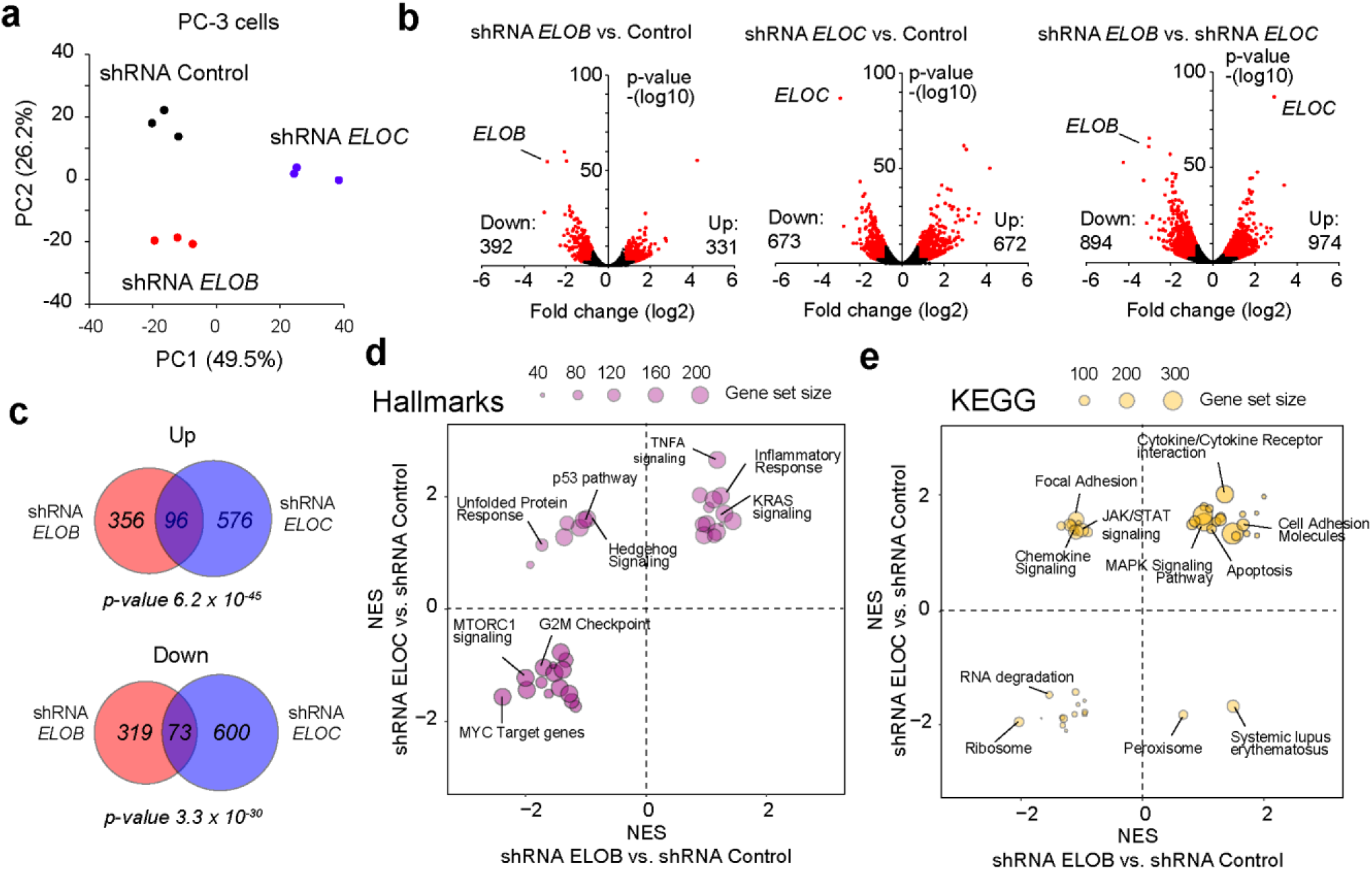
Knockdown of *ELOB* and *ELOC* strongly influences the cellular program of PC3 prostate cancer cells. **a**, PCA (Principal Component Analysis) of the RNA-Seq data upon knockdown of *ELOB* and *ELOC*. **b**, Volcano plots showing the gene expression changes between the three investigated conditions. Red indicates significantly differentially expressed genes (*p* < 0.01, log2 fold change > 0.75). **c**, Overlap of upregulated and downregulated genes in *ELOB* and *ELOC* knockdown cells. The *p*-values were calculated via a hypergeometric probability test. **d**, Comparison of dysregulated gene sets after *ELOB* and *ELOC* knockdown, determined via GSEA (gene set enrichment analysis)^21^. The hallmark and KEGG gene sets are based on MSigDB.

Nonetheless, gene set enrichment analysis (GSEA) suggested that many gene sets related to cell growth, such as MYC target genes, the G2M checkpoint, as well as signalling pathways, such as KRAS and MAPK, were similarly affected upon knockdown of *ELOB* and *ELOC* (**Fig. 2d**). Other pathways, such as the p53 pathway and hedgehog signalling, were distinctly affected by the two knockdowns (**Fig. 2d**). Thus, these results suggest that *ELOB* and *ELOC* are essential for cancer cell growth by maintaining key cellular pathways, but they may have non-identical functions.

Together we conclude that ELOB and ELOC play an important function in cancer cell growth, making them attractive targets for cancer therapy.

### EPOP’s BC-box does not require other proteins for interaction with ELOB/C

The main function of ELOB and ELOC proteins has been attributed to the interaction of the ELOB/C heterodimer with BC-box containing proteins^7,11,12^. We therefore hypothesised that blocking the interaction between ELOB/C and BC-box proteins could be a valid option to interfere with the function of these proteins and inhibit cancer cell growth.

Importantly, most proteins that interact with ELOB/C also possess a cullin-box, in addition to the BC-box^7^ (**Extended Data Fig. 2a**). This combination allows the formation of multiprotein complexes, involved in protein ubiquitination^7^. Indeed, previously published mass spectrometry data^15^ that investigated the interactome of ELOB showed that it interacts with cullin 2 and cullin 5 and with many proteins that possess both a BC-box and a cullin-box (**Extended Data Fig. Fig 2b**). However, not all ELOB/C binding proteins possess a cullin-box. One major exception is EPOP, which is a PRC2 (Polycomb repressive complex 2) interacting protein^14,15^. EPOP does not possess an obvious cullin-box, and previous mass spectrometry data^14,15^ do not support that EPOP interacts with cullins. Thus, we concluded that EPOP likely does not require cullins for its interaction with ELOB/C.

However, to date it has not been addressed whether EPOP may require other proteins for the interaction with ELOB/C. To assess this question, we immunoprecipitated EPOP from HEK293 cells, followed by Western blot and mass spectrometry analysis. First, we confirmed that mutation of the BC-box (L40A) abrogates the interaction with ELOB but not with the PRC2 components SUZ12 and EZH2 (**Fig. 3a**). Oppositely, deletion of the C-terminal region of EPOP, which is important for the interaction with PRC2^22^, prevents interaction with SUZ12 but not with ELOB and ELOC (**Fig. 3a**), suggesting that EPOP’s interaction with PRC2 and ELOB/C are independent of each other. Via unbiased semiquantitative mass spectrometry, we validated that EPOP interacts with PRC2 and ELOB/C (**Fig. 3b**). This experiment also confirmed previous reports that EPOP predominantly interacts with the PRC2.1 class of PRC2, which consists of the Polycomb-like proteins, but lacks JARID2 and AEBP2^14-16^. Importantly, we found that the L40A mutation of the BC-box exclusively impaired the interaction with ELOB/C, while no other proteins were affected by this mutation (**Fig. 3c**). Deletion of the C-terminus of EPOP (ΔCTR) abolished the interaction with PRC2 but only mildly affected the interaction with ELOB/C (**Fig. 3d**), confirming that the interaction of EPOP with PRC2 is not essential for EPOP’s interaction with ELOB/C. Together, these results suggest that EPOP does not appear to require other proteins for the interaction with ELOB/C. We therefore conclude that the BC-box of EPOP is not only required but also sufficient for the interaction with ELOB/C. Consequently, we hypothesised that EPOP’s box likely has a high binding affinity to ELOB/C, allowing an efficient association with ELOB/C without additional interaction partners.

**Fig. 3:**
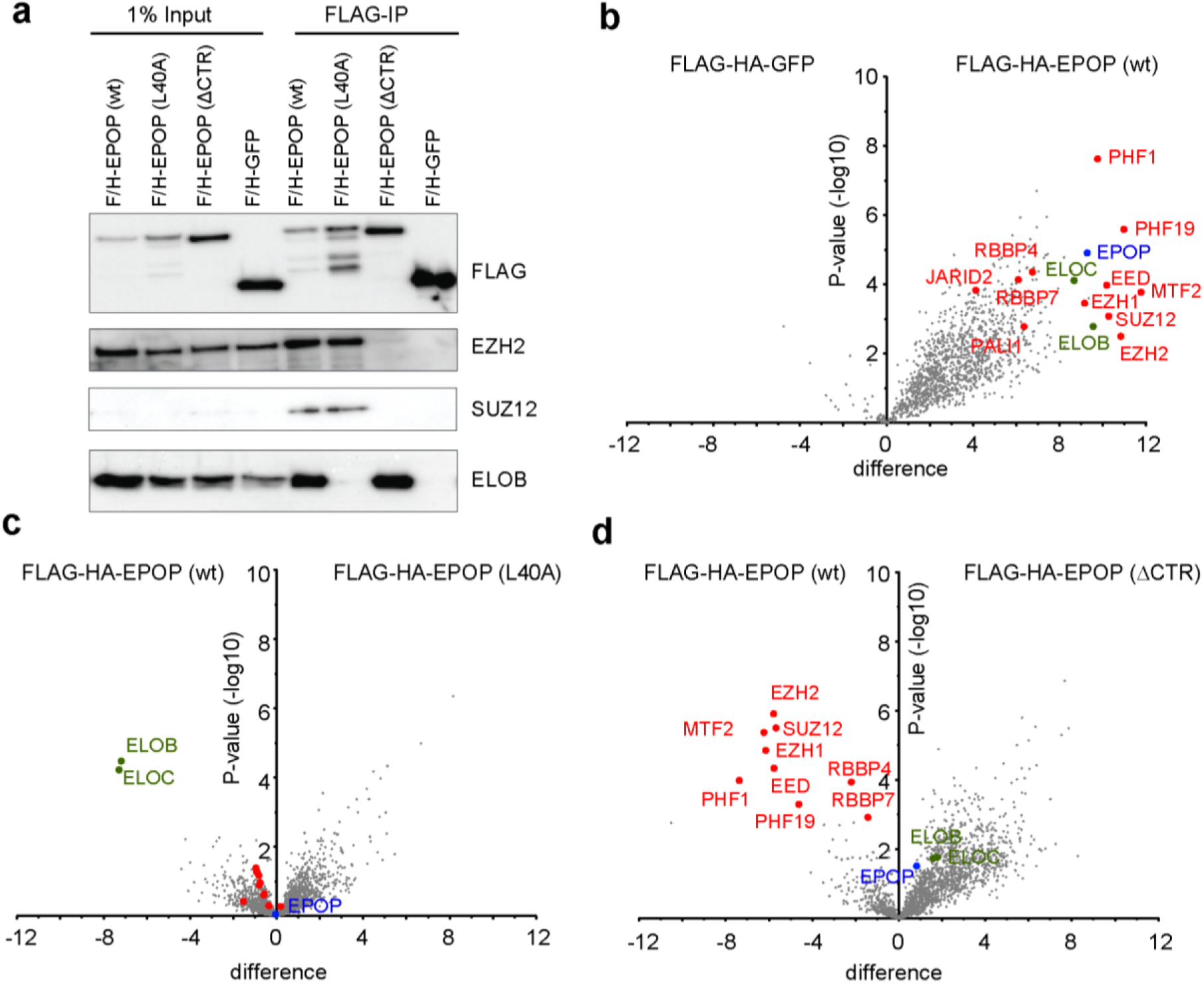
EPOP interacts with ELOB/C independent of other proteins. **a**, Western blot of semi-endogenous co-immunoprecipitation experiments using wild-type, L40A mutated and C-terminal truncated FLAG-HA-(F/H)-tagged EPOP as bait. The experiment was performed in HEK293 cells. **b-d**, Semi-quantitative mass spectrometry upon co-immunoprecipitation of various FLAG-HA-EPOP constructs presented as Volcano plot, reflecting four biological replicates. EPOP is marked blue, ELOB and ELOC are green and PRC2 components are marked red. **b**, Comparison of FLAG-HA-EPOP wild-type (w int) versus FLAG-HA-GFP control. **c**, Comparison of FLAG-HA-EPOP (L40A) versus FLAG-HA-EPOP (wt). **d**, Comparison of FLAG-HA-EPOP (ΔCTR) versus FLAG-HA-EPOP (wt). Silver staining of the co-immunoprecipitated material for b-d are presented in Supplementary Fig. S20.

### The EPOP BC-box specifically binds to Elongin BC with subnanomolar affinity

Thus, we selected the BC-box of EPOP to explore Elongin BC inhibition. We synthesised a fluorescently labelled EPOP BC-box peptide (EPOP_wt_-FAM, **Fig. 4a**), where the 5(6)-carboxyfluorescein was orthogonally introduced in a C-terminal lysine separated by two units of 6- aminohexanoic acid (Ahx) to avoid binding interferences. The interaction of this peptide with ELOB/C was studied by fluorescence polarization (FP), and compared with the previously reported fluorescently labelled BC-box of HIV-Vif (FAM-HIV-Vif, **Fig. 4a**)^23^. The observed inconsistencies in the dissociation constants (*k*_D_) suggested to us that both peptides are tight binding inhibitors^24^, and therefore the concentration of the tracer must be decreased to avoid overestimation of the *k*_D_ (**Extended Data Fig. 4a**). Gratifyingly, our saturation experiments demonstrated a direct interaction between EPOP_wt_-FAM and the heterodimer Elongin BC. Although the dynamic range is larger for FAM-HIV-Vif, our EPOP-derivative peptide displayed stronger affinity than FAM-HIV-Vif, reaching subnanomolar affinities (*k*_D_ = 0.45 ± 0.03 nM versus *k*_D_ = 3.52 ± 0.33 nM, **Fig. 4b**).

**Fig. 4:**
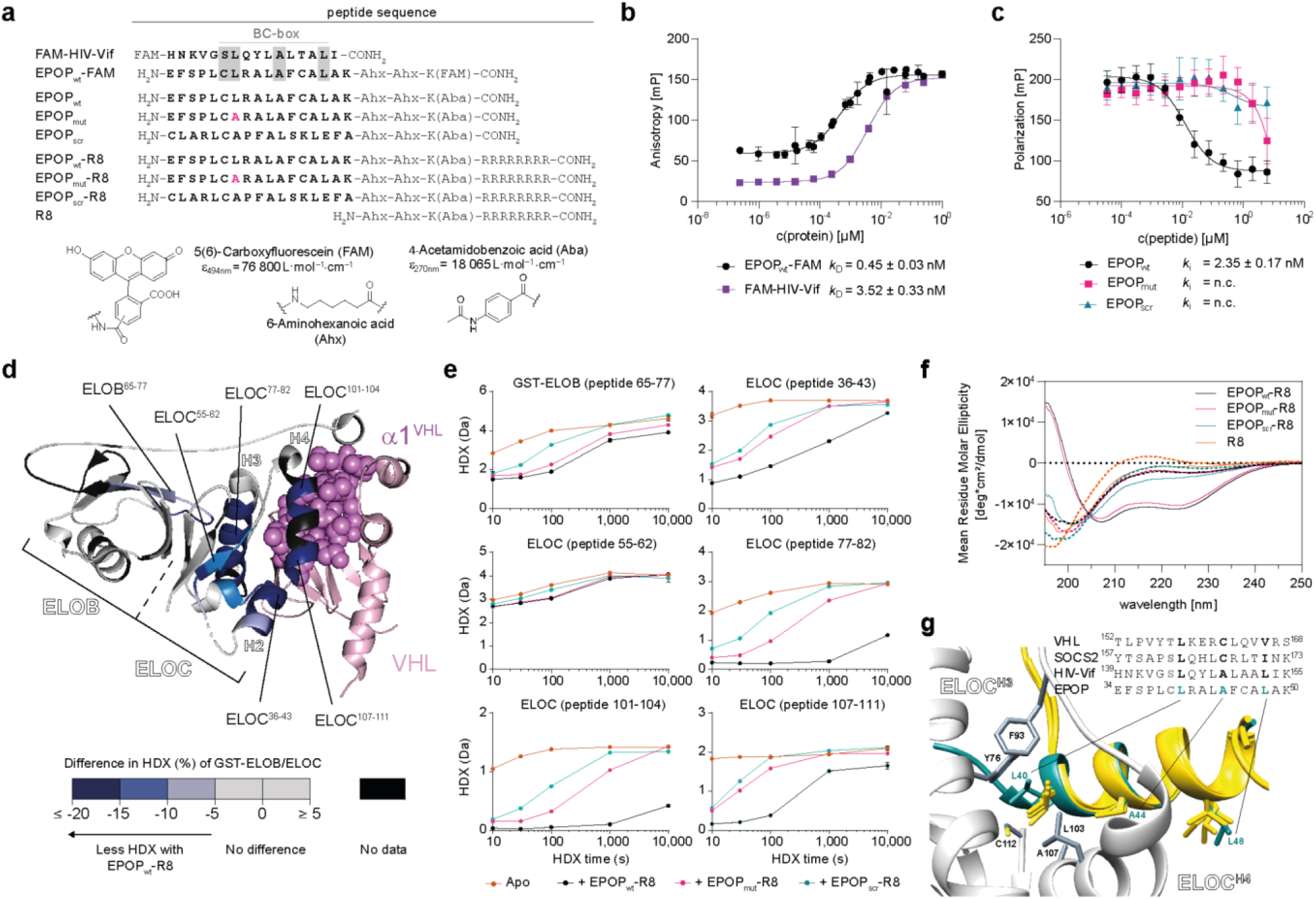
Synthetic EPOP BC-box helix recognises Elongin BC and binds with subnanomolar affinity. **a**, Synthetic BC-boxes derived from both EPOP and reported HIV-Vif binder. EPOP sequences were modified with an aminohexanoic acid-based linker, bearing either a chromophore (Aba) or fluorophore (FAM), and an R8 cell-penetrating sequence. **b**, Fluorescence polarization-based binding experiments of EPOP_wt_-FAM and FAM-HIV-Vif (0.1 nM) to GST-ELOB/ELOC in 25 mM Tris-Cl pH 7.5, 100 mM NaCl, 0.1 mM TCEP and 0.02% Triton X-100. **c**, Competitive fluorescence polarization assay in the presence of 0.1 nM of the complex EPOP_wt_-FAM with GST-ELOB/ELOC under the same conditions as in b. **d**, HDX difference in the GST-ELOB/ELOC complex with(out) EPOP-derivatives peptides mapped on HIF-1α/VHL/ELOB/ELOC crystal structure (PDB 1LM8)^25^. Peptide-dependent reduction in HDX clusters along the α1^VHL^ (pink spheres, residues 155-170) binding site of ELOC. **e**, Representative GST-ELOB and ELOC peptides (residue numbers in brackets) displaying differences in HDX depending on EPOP_wt_-R8, EPOP_mut_-R8 and EPOP_scrt_-R8. Data represent the mean ± s.d. of n=3 technical replicates. **f**, Circular dichroism spectra of EPOP-derivative peptides at 25 μM in 5 mM Tris-Cl pH 7.5 in the absence (dotted lines) or presence (solid lines) of 25 mM sodium lauryl sulfate. **g**, Structural overlay of the predicted EPOP^43-50^ (turkey) by AlphaFold with pVHL (1LM8), SOCS2 (2C9W) and HIV-Vif (3DCG)(yellow)^23,25,26^.

Next, we studied interaction specificity. For this purpose, we synthesised three additional peptides: a wild-type variant (EPOP_wt_), a scramble one (EPOP_scr_), and a singly mutated wild-type derivative, in which the leucine 40 was exchanged by an alanine (EPOP_mut_) (**Fig. 4a**). This mutation has been reported to interrupt the interaction of EPOP with ELOB/C^14^, which we reconfirmed by our mass spectrometry experiments (**Fig. 3c**). The *p*-aminobenzoic acid (Aba) replaced the fluorophore to ensure accurate concentration determination with minimal structural change. Once the peptides were prepared, their binding affinity was evaluated by competitive FP-based experiments using EPOP_wt_-FAM as a tracer. As expected, the exchange of the chromophore hardly affects the interaction (**Fig. 4c**). More importantly, only the wild-type variant can efficiently displace the tracer with an apparent inhibitory constant (*k*_i_) of 2.35 ± 0.17 nM. However, the apparent *k*_i_ for both EPOP_mut_ and EPOP_scr_ could not be calculated at the applied concentrations. These measurements corroborated that the replacement of the L40 resulted in a strong loss of interaction (*k*_i_ of μM versus nM). Both the specificity and the high affinity of EPOP_wt_ to Elongin BC encouraged us to investigate whether this peptide can indeed affect Elongin BC in living cells and, consequently, its related functions. To this end, analogue peptides carrying the octaarginine as a vector (EPOP_wt_-R8, EPOP_mut_-R8 and EPOP_scr_-R8) as well as the cell-penetrating peptide alone (R8) were synthesised and further investigated.

### Synthetic EPOP BC-box binds to the H3-H4 groove of ELOC

To experimentally probe whether the synthetic EPOP BC-box peptide would bind to ELOB/C in the same way as previously reported BC-box peptides^9,23,25,26^, we performed hydrogen/deuterium exchange (HDX) mass spectrometry on the complex of ELOB fused N-terminally to glutathione *S*-transferase (GST) and ELOC (GST-ELOB/ELOC complex). The HDX of the GST-ELOB/ELOC complex in the presence of EPOP peptides was compared with that of apo-GST-ELOB/ELOC, whereby differences in HDX would reflect alterations in the GST-ELOB/ELOC conformation in dependence of EPOP peptides (**Fig. 4d-e, Supplementary Figs. S12-S14**). Reduced HDX of ELOC in the presence of EPOP_wt_-R8 was particularly observed for the peptides clustering along H2-H4 of ELOC (**Fig. 4d-e**, ELOC^36-43^, ELOC^77-82^, ELOC^101-104^ and ELOC^107-111^), correlating well with the structurally validated binding site of the BC-box containing helix α1 of VHL in the HIF-1α/VHL/ELOB/ELOC complex (PDB 1LM8)^25^. Quite unexpectedly, a decrease in HDX was also apparent for residues 63-70 of ELOB (**Fig. 4e**, representative peptide ELOB^65-77^, **Extended Data Fig. 4b**), which may indicate a broader impact of EPOP peptide-binding on the topology of the ELOB/ELOC complex beyond the BC-box binding site at ELOC. The EPOP_mut_-R8 and EPOP_scr_-R8 peptides induced perturbations in HDX in similar areas of ELOC (**Fig. 4e, Supplementary Figs. S12-S14**), albeit at lower amplitudes, reflecting their different binding affinities (**Fig. 4c**). This finding highlights the critical role of EPOP^L40^ (**Fig. 4f**) in establishing the interaction between EPOP and ELOC.

To rule out different secondary structures among peptides, we conducted circular dichroism spectroscopy (**Fig. 4f**). As expected from the high sequence analogy, EPOP_wt_-R8 as well as EPOP_mut_-R8 underwent a comparable increase in their helical content under simulated hydrophobic conditions with SDS, (from 6% to 27% for EPOP_wt_-R8 versus from 7% to 29% for EPOP_mut_-R8) while EPOP_scr_-R8 and R8 did not change. In our attempt to obtain a molecular interpretation, we compared the predicted structure (EPOP_sim_) to the reported α1-VHL, SOCS2 and HIV-Vif in complexes with Elongin BC (**Fig. 4g**). On the one hand, the major contribution of EPOP^L40^ to Elongin BC recognition is evident because of the most severe local H/D exchange decrease in the ELOC^77-82^ and ELOC^101-104^ regions (**Fig. 4e, Supplementary Figs. S12-13)**. Indeed, as described by Stebbins *et al*.^9^, our data confirmed that ELOC^Y76^, ELOC^L102^ and ELOC^A107^ are the main contributors within the ELOC hydrophobic pocket. No alternation was observed in the ELOC^87-98^ region, suggesting that it does not contribute to the stabilisation of EPOP^L40^ (**Supplementary Figs. S12-14**). Of note, all BC-box peptides adopt an α-helical conformation parallel to the C-terminal helix 4 (H4) of ELOC, where the conserved leucine residue (EPOP^L40^ VHL^L158^, SOCS2^L163^ and HIV-Vif^L145^) is embedded into the ELOC hydrophobic pocket. According to our simulations, EPOP_sim_ seems to have ideal interactions, which are only partially fulfilled by its homologues (**Extended Data Fig. 4c-f)**. In particular, the presence of EPOP^F45^ should provide a stronger Van-der-Waals interaction with ELOC^L104^ than with the analogue VHL^L163^ or HIV-Vif^L150^. Along these lines, EPOP^R41^ additionally may form a stronger salt bridge with ELOC^N108^ than in the other BC-boxes with VHL^K159^. Altogether, our structural data justify the high affinity of EPOP_wt_ recognition by ELOB/C.

### Interaction of ELOB/C with EPOP and VHL is inhibited by BC-box peptides

To address whether the BC-box peptides inhibit the protein interaction of ELOB/C, we performed pull-down experiments in vitro. Immobilised GST-ELOC/ELOB were incubated with cellular extracts expressing FLAG-tagged EPOP. Consistent with previous results^14^, EPOP can be pulled-down by GST-ELOC/ELOB, but not with GST alone (**Fig. 5a, b**, left two lanes). Subsequently, we investigated the consequence of the presence of the BC-box peptides during this experiment. We found that at a concentration of 20 μM the wild-type BC-box peptide, but not the mutated or scrambled peptide, prevented EPOP from being pulled-down with GST-ELOC/ELOB (**Fig. 5a**). This inhibitory effect occurs in a dose-dependent manner (**Fig. 5b**), thus suggesting that the wild-type BC-box peptide efficiently prevents the interaction of EPOP with ELOB/C.

**Fig. 5:**
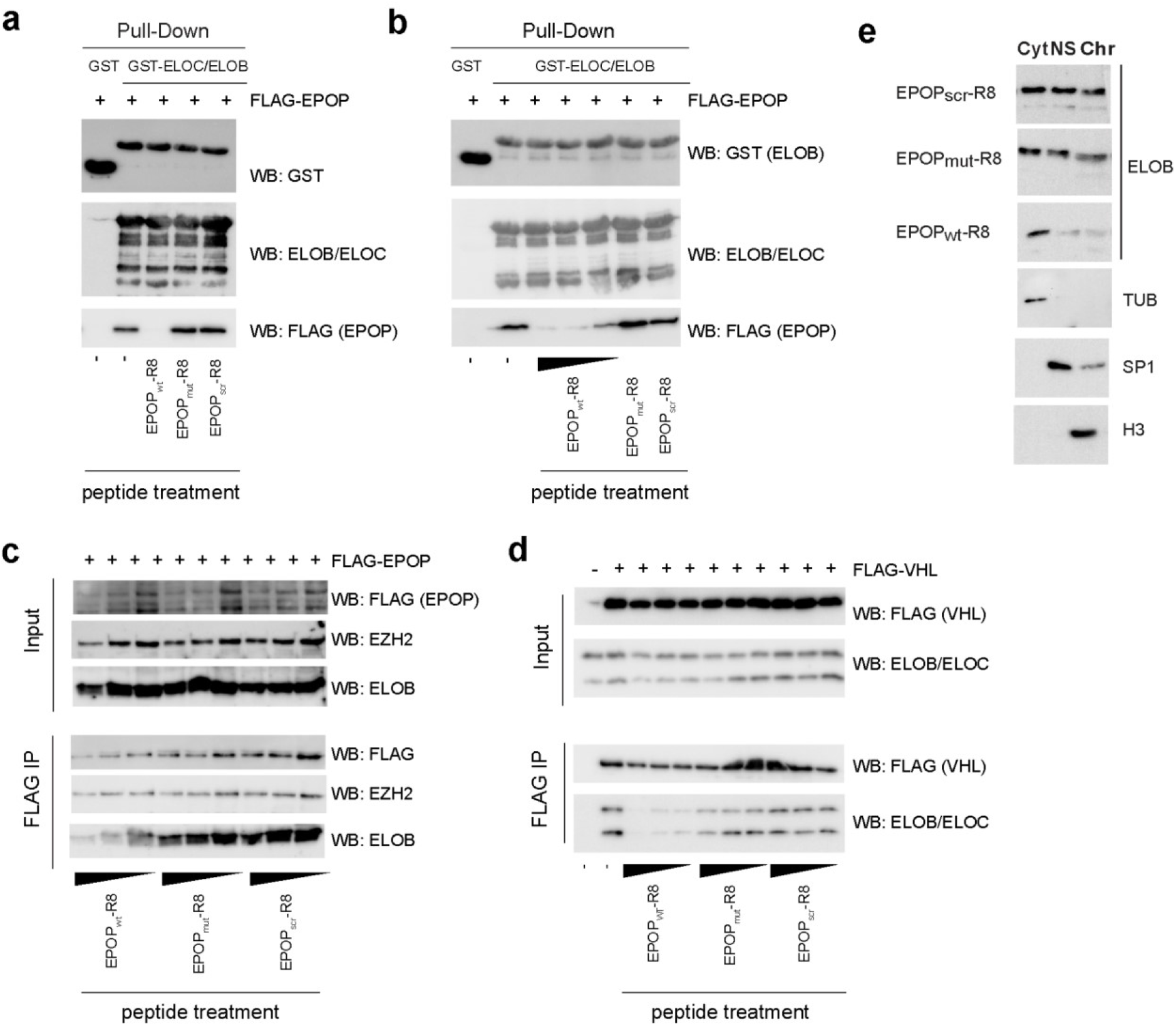
EPOP BC-box peptide inhibits the interaction of ELOB/C with its partners. **a**, GST-pulldown using GST-ELOC/B as bait. The interaction of EPOP with GST-ELOC/B is inhibited by the presence of EPOP_wt_-R8 peptides (20 μM), but not with mutated or scrambled peptides. **b**, As in a, but with increasing amounts of EPOP_wt_-R8 peptide (6.25, 12.5, 20 μM). **c**, Western blot of semi-endogenous co-immunoprecipitation experiment with FLAG-EPOP as bait. The interaction with ELOB but not the PRC2 member EZH2 is reduced by the presence of the EPOP_wt_-R8 peptide (12.5 μM). **d**, As in c, but using FLAG-VHL as bait. For a-d, one representative result of at least 2 biological replicates is shown. Additional replicates of c and d are presented in Supplementary Fig. S21. **e**, Representative ELOB Western blot after cellular fractionation of PC3 cells upon peptide treatment. Cyt = cytoplasm, NS = nucleoplasm, Chr = chromatin.

To address whether this interaction is also inhibited in a cellular context, we performed co-immunoprecipitation experiments with ectopically expressed FLAG-tagged EPOP in HEK293 cells. In presence of the EPOP_mut_-R8 or EPOP_scr_-R8 peptide at a concentration of 12.5 μM, we could efficiently co-immunoprecipitate ELOB together with FLAG-EPOP (**Fig. 5c**). In contrast, an increasing amount of the EPOP_wt_-R8 peptide reduces the level of co-immunoprecipitated ELOB. This reduced co-immunoprecipitation can likely be attributed to the inhibition of the interaction of EPOP with ELOB/C, consistent with our in vitro pull-down experiments. In contrast, the interaction of EPOP with the PRC2 core member EZH2 was not strongly affected upon the peptide treatment (**Fig. 5c**), suggesting that the peptide does not interfere with the interaction between EPOP and PRC2. This result is consistent with our mass spectrometry results, which showed that EPOP interacts with ELOB/C and PRC2 independently (**Fig. 3**).

To validate the inhibitory function of the BC-box peptide for another ELOB/C-BC-box interaction, we performed similar experiments with the ELOB/C interaction partner VHL^11^. Affirmatively, we found that the addition of the EPOP_wt_-R8 but not the control peptides prevented the co-immunoprecipitation of ELOB and ELOC (**Fig. 5d**). In line with an impaired interaction of ELOB/C with nuclear interaction partners, such as EPOP and VHL, we observe a reduced level of ELOB in the nuclear and chromatin fraction upon treatment with the wild-type peptide (**Fig. 5e**). Thus, these experiments suggest that the EPOP_wt_-R8 peptide can block ELOB/C from interacting with its interaction partners in cells, while the EPOP_mut_-R8 peptide cannot.

### Peptide treatment in cancer cell lines reduces cell proliferation both in clonogenicity and via apoptosis induction

Next, we explored whether the specific interference of our synthetic EPOP-derivative peptides with Elongin BC could be translated into an inhibition of cancer cell proliferation. We first conducted cell viability assays in PC3, MCF-7, and SH-SY5Y cells. While both PC3 and MCF-7 cell viability were analysed via a resazurin-based assay, SH-SY5Y cells were studied by an ATP-dependent luciferase-based assay due to the low metabolic throughput of SH-SY5Y (**Supplementary Fig. S15**), which required a high sensitivity assay. All cancer cell lines manifested dose-response impaired growth by the addition of EPOP_wt_-R8 over 24 h (**Fig. 5a, Supplementary Fig. S16)**. This effect was statistically significantly different in comparison to EPOP_mut_-R8 and EPOP_scr_-R8 in all cell lines (**Fig. 5b)**. The highest differences were observed for SH-SY5Y (3-fold cell growth inhibition between EPOP_wt_-R8 and EPOP_scr_-R8 at 21.6 μM) (**Fig. 5b**); however, the strongest cancer growth inhibition was observed in PC3. Importantly, the cell-penetrating sequence (R8) did not affect viability (**Supplementary Fig. S17**). These experiments, together with our in vitro studies, strongly suggest the sequence dependency of our EPOP-derived peptides in living cells. Due to these observations, we investigated PC3 cells in colony formation assays (**Fig. 5c,d**). We observed that treating the cells with the EPOP_wt_-R8 peptide, but not with the control peptides, led to significantly reduced colony formation, further supporting that inhibiting ELOB/C prevents optimal cell growth.

To understand the molecular reasons, we investigated the consequence of ELOB/C inhibition via the BC-box peptide on the transcriptional landscape. We treated the cells with EPOP_wt_-R8, EPOP_mut_-R8 and EPOP_scr_-R8 peptides and performed RNA-Seq experiments after 24 hours of treatment in three replicates. Principal component analysis demonstrated that the EPOP_wt_-R8, and EPOP_scr_-R8 treated cells differed in their transcriptome, while the cells treated with the EPOP_mut_-R8 peptide were between the wild-type and the scrambled versions (**Fig. 6e**). This observation is consistent with our previous results, indicating that the EPOP_mut_-R8 peptide has some residual activity (**Fig. 6a**). For further investigation, we focused on the differences between cells treated with EPOP_wt_-R8 and EPOP_scr_-R8 peptide. We identified approximately 100 genes that were differentially expressed (log2-fold change > 0.35, *p*-value < 0.01) (**Fig. 6f**), indicating that the impact of the peptide treatment is subtler than that of the knockdown of ELOB or ELOC. GSEA showed that a substantial number of pathways were significantly dysregulated, supporting a shift in the transcriptional landscape in the EPOP_wt_-R8 treated cells, compared to control cells. Looking at cancer pathways in general, we confirmed that inhibiting ELOB/C with our peptide inhibitor leads to the downregulation of pathways in cancer (**Fig. 6g**). Closer investigations of the dysregulated pathways showed an upregulation of the G2/M checkpoint, sister chromatin segregation, and cell division (**Extended Data Fig. 5a**), implicating a dysregulation of cell cycle related genes. Indeed, we found that many genes related to the cell cycle were upregulated (**Extended Data Fig. 5b)**, consistent with an altered cell cycle progression of the EPOP_wt_-R8 treated cells compared to the scrambled control (**Extended Data Fig. 5c, Supplementary Fig. S22**). Oppositely, we observed a downregulation of many signalling pathways including the TNF alpha/NF-kappa, cytokine JAK/STAT and the hypoxia signalling pathways (**Extended Data Fig. 6a**). Consistently, the genes related to epithelial mesenchymal transition (EMT), which strongly depend on signalling pathways^27^, were also significantly downregulated (**Extended Data Fig. 6b**). The changes in the signalling pathways could be due to altered function of the SOCS proteins, a group of common interaction partners of ELOB/C^7,10^. The effect on hypoxia-related pathways may be linked to the role of ELOB/C for the regulation of VHL^9^, which is an important player in the hypoxia response^28^. This hypothesis is supported by the observation that the genes of *ELOB*, but not *ELOC*, and *VHL* positively correlate in CRISPR-screening experiments (**Extended Data Fig. 6c, d**), suggesting that they may work together in several cancer types. However, given the versatility of target proteins of ELOB/C, both in the cytoplasm and in the nucleus, it is likely that a large number of distinct cellular processes are influenced by inhibiting ELOB/C. A closer inspection of the molecular processes influenced by ELOB/C may provide a clearer picture of how inhibiting ELOB/C affects cellular functions. Unexpectedly, the gene expression changes upon peptide treatment showed almost no correlation with the effects observed after the knockdown of *ELOB* or *ELOC* (**Extended Data Fig. 6e, f**). One possible reason could be that the gene expression changes of the peptide treatment were measured after a shorter time period (24 h), compared to the knockdown cells (4 days), which may lead to less severe effects. Another possibility is that the peptide treatment interferes only with the function of the ELOB/C dimers, while the knockdown may additionally affect ELOB/C dimer-unrelated functions of the ELOB and ELOC proteins (see discussion).

**Fig. 6:**
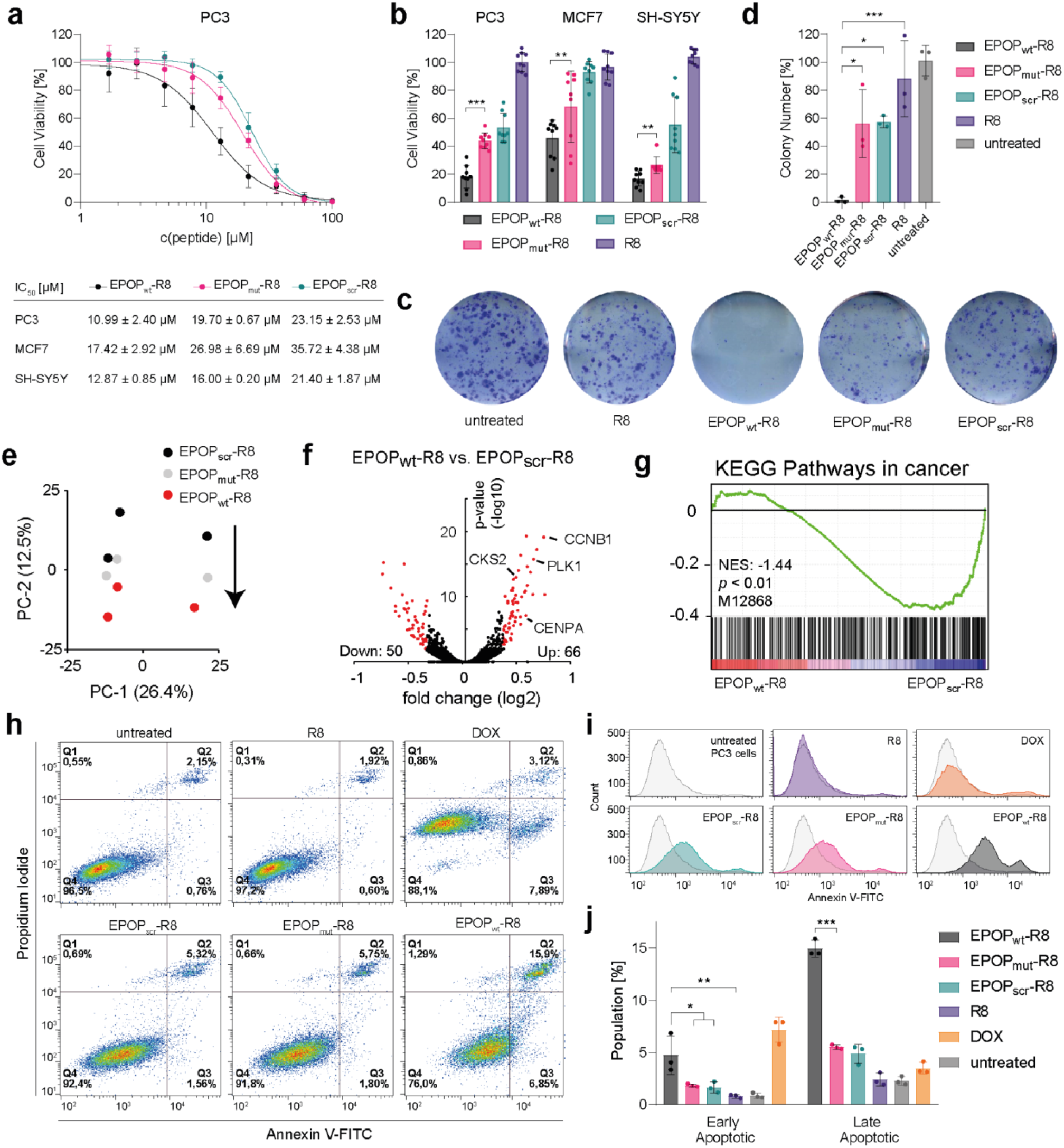
EPOP BC-box peptide inhibition interferes with cell proliferation, colony-forming capability, and cancer-related pathways and induces apoptosis in PC3 cells. **a**, Dose-response cell viability in PC3 cells and IC_50_ values for all tested cancer cells after 24 h of incubation. Data in the dose-response curve represent the mean ± s.d. of n = 3 experimental independent replicates (three technical replicates each). **b**, Cell viability after treatment with 21.6 μM EPOP-derivative peptide. Data represent the mean ± s.d. of n = 3 experimental independent replicates (three technical replicates each). Significance was evaluated by a two-tailed unpaired Student’s t-test. **c**, Representative results of PC3 cells stained with methylene blue upon colony formation assay in presence of EPOP_wt_-R8, and controls. **d**, Quantification of colony formation assay. Cells were treated for 24 h with 6.5 μM of the respective peptides and grown for 10 days before analysis. Data represent the mean ± s.d. of three biological replicates. Significance was evaluated by a two-tailed unpaired Student’s t-test. **e**, PCA analysis of RNA-Seq data after peptide inhibition. The arrow indicates the shift gene regulatory network upon treatment with the wildtype peptide. **f**, Volcano plot of the RNA-Seq data showing the significantly differentially expressed genes (log2-fold change > 0.35; *p*-value < 0.01). **g**, GSEA analysis showing the downregulation of pathways in cancer. **h**, Flow cytometry experiments of PC3 cells treated with 5 μM EPOP BC-box peptides over 48 h under the conditions in a, to determine apoptosis induction via Annexin V-FITC and propidium iodide staining. A representative experiment is shown here, replicates and gating strategy are shown in the supplementary information (**Supplementary Fig. S18, S19**). **i**, Histograms of Annexin V-FITC stained PC3 cells shown in h. **j**, Statistical analysis of Annexin V-FITC/PI stained cell samples. Data represent mean ± s.d. of n = 3 experimental independent replicates. Significance was evaluated by a two-tailed unpaired Student’s t-test. *, *p* < 0.05; **, *p* < 0.01; ***, *p* < 0.001.

Considering the changes in cellular pathways as well as reduced cell viability, we hypothesised that apoptosis induction may be the underlying cause. To this end, we conducted flow cytometry experiments with prostate cancer (PC3) cells, exposed to 5 μM of the corresponding EPOP BC-box peptides over 48 h (**Fig. 5h**) and compared the effects with the reported apoptotic doxorubicin (DOX) at 1 μM. DOX caused a slight increase in the propidium iodide (PI) signal, which was compensated by suitable quadrant placing, and consequently yielding comparable results to the literature^29,30^. While R8-treated cells were indistinguishable from untreated cells, our data showed that EPOP_wt_-R8 remarkably increased early apoptotic cells by a 4-fold compared to EPOP_mut_-R8 and EPOP_scr_-R8 (Q3, 6.85% versus 1.80% and 1.56%, respectively) (**Fig. 5h**). Furthermore, an almost 3-fold difference in late apoptotic cells could be observed (Q2, 15.9% versus 5.75% and 5.32%, respectively). In contrast to DOX, which foremost increased early apoptotic cells (7.89%), EPOP_wt_-R8 predominantly induced late apoptotic cells. Besides, the corresponding histograms clearly illustrated the increment of the population shift to apoptotic-positive populations after peptide treatment (**Fig. 5i**). Of note, our three independent replicates demonstrated a statistical significance between EPOP_wt_-R8 and the L40A mutant EPOP_mut_-R8 (**Fig. 5j, Supplementary Fig. S18**). Altogether, our results suggest that blocking the function of ELOB/C leads to reduced cell proliferation, which may be at least in part due to the induction of apoptosis.

## Discussion

Aberrant functions of multiprotein complexes often lead to the development of human diseases, such as cancer^1^. The functionality of the proteins is typically altered either due to mutations or dysregulated expression^31^. The identification of key elements that are involved in disease development and progression is the first step in drug discovery. Here, we identified the heterodimer ELOB/C as a potential drug target for cancer. We demonstrate that the protein binding function of ELOB/C can be inhibited via a peptide, based on the ELOB/C-binding motif of EPOP.

Both ELOB and ELOC belong to the group of genes that are designated as “common essential” in most cancer types (**Fig. 1**)^18^, thus playing a crucial role in cancer cell proliferation. However, to date no efforts have been made to develop inhibitors for these proteins. Given that the main function of the heterodimeric ELOB/C has been linked to the binding of BC-box containing proteins^7,12,14,26^, we developed a peptide inhibitor that blocks this interaction. We used the ELOB/C-binding sequence of EPOP and demonstrated that EPOP’s BC-box sequence alone is sufficient for ELOB/C recognition. Our newly developed peptide binds to the BC-box recognition pocket of ELOB/C with a binding affinity in the nanomolar range (**Fig. 4c**). Our peptide surpasses the binding affinity of other BC-boxes in vitro (**Fig. 4b**) supporting the effectiveness of our strategy. With future studies, our proof-of-concept inhibition in cells opens the possibility of exploring alternative strategies such as stapled peptides^32^, peptidomimetics and small molecules to inhibit ELOB/C^3^.

Importantly, until now ELOB and ELOC were mostly considered as functional tandem^7^, and to the best of our knowledge, no individual roles have been described. However, the existence of independent functions of ELOB and ELOC is supported by several observations. First, the consequences of ELOB and ELOC knockouts in cancer cells did not correlate well (**Extended Data Fig. 1a**). Second, the cellular localization of ELOB and ELOC only partially overlapped (**Extended Data Fig. 2**), suggesting that they can exist in the cells as monomers or in complex with other proteins. Third, our knockdown experiments showed that the depletion of ELOB and ELOC had different consequences on gene expression (**Fig. 1c**). These findings suggest cellular roles of ELOB and ELOC that lie beyond the ELOB/C dimer. These additional functions may explain why inhibiting ELOB/C with our peptide inhibitor (**Fig. 6f**) has less severe consequences compared to the knockdown of ELOB and ELOC (**Fig. 2b**). Thus, we currently cannot rule out the possibility that inhibiting the ELOB/C dimer using our peptide inhibitor interferes only with the functions that are related to the BC-box binding function of ELOB/C, while other functions of ELOB and ELOC may be unaffected. These other functions may also be of relevance for cancer cell growth. Further experiments will be essential to obtain a clear picture of which biological functions are dependent on the ELOB/C dimer, hence can be inhibited by our peptide, and which functions of ELOB and ELOC are facilitated independently of each other.

Nonetheless, application of the inhibiting peptide in cancer cells showed consistent inhibition of cancer cell proliferation throughout the tested cell lines (**Fig. 6b**). Consolidating conclusions were found in colony formation assays and flow cytometry-based apoptosis assays (**Fig. 6c-g**), all pointing towards compromised viability functions of cancer cells upon treatment with the peptide inhibitor. Thus, our peptide inhibitor provides a versatile tool to perform further mechanistic studies and to identify the cellular pathways that are dependent on the ELOB/C dimer. This work would provide insights why cancer cells require ELOB and ELOC for cell growth and whether ELOB/C blockage could contribute to cancer treatment.

## Methods

### Solid-phase peptide synthesis

All peptides were synthesized according to the standard Fmoc-SPPS methodology. Tentagel TG S RAM resin was swollen 30 min prior to synthesis. The synthesis was performed with either the INTAVIS ResPep SLi (5 μmol scale) or Advanced ChemTech Apex 396 (20 μmol scale) peptide synthesizer. Fmoc-protected N-termini were removed with 20% piperidine in DMF. Fmoc-amino acids (5.3 eq) were coupled with DIC (5.3 eq) and Oxyma (5.3 eq) at 0.24 M in 110 μL DMF (9.0 eq at 0.25 M in 720 μL for Apex 396). The K/A-Ahx-Ahx-K(Alloc) sequence was previously synthesized by hand using HATU as coupling reagent (conditions: 4 eq Fmoc-amio acid, 4 eq HATU and 8 eq DIEA at 0.5 M in 160 μL DMF). For the acetylation step 2,5-lutidine (6%) and Ac_2_O (5%) in 500 μL DMF were used. Orthogonal lysine side-chain deprotection was performed by incubating the resin with Pd(OAc)_2_ (0.2 eq), PPh_3_ (3 eq), PhSiH_3_ (10 eq) and NMM (10 eq) in 2 mL degassed CH_2_Cl_2_ for 1 h^33,34^. Subsequently either 4-acetamidobenzoic acid (10 eq) was coupled with HATU/DIEA at 0.5 M in DMF or 5(6)-carboxyfluorescein (20 eq) with DIC/Oxyma at 0.5 M in 800 μL DMF. Final peptides were cleaved with 2 mL of 82.5% TFA, 5% phenol, 5% H_2_O, 5% thioanisole and 2.5% EDT (1.5x volume for R8 containing peptides). After precipitation with diethyl ether, the obtained peptides were purified by preparative RP-HPLC on an Agilent Infinity II 1260 system. The columns used for the purification and final characterization are provided in the Supplementary Table S1. High-resolution mass spectrometry (HR-ESI-MS) was performed using a TQ-FT Ultra mass spectrometer (Thermo Fisher Scientific). The characterization of the peptides used in this study is shown in Supplementary Figs. S1-11.

### UV-vis spectroscopy

Concentration determination was performed on a Tecan Spark 20M multimode microplate reader at room temperature. All measurements were performed in a 1.4 mL quartz cuvette (Hellma Analytics 104F-QS) with a pathlength of 1 cm. Increasing amounts of the to-be-measured peptide were added to an 800 μL solution of the respective blank, maintaining the increase in the added volume below 10% of the initial solution. Absorbance spectra were recorded, and the peptide concentrations were determined using the following extinction coefficients: fluorescently tagged peptides, in 0.1 M PBS adjusted to pH 9.0 ε_494nm_ = 76,900 L*mol^-1^*cm^-1^, Aba-modified peptides as DMSO stocks were determined in ultrapure water ε_270nm_ = 18,069 L*mol^-1^*cm^-1^)^35-37^.

### Circular dichroism spectroscopy

25 μM concentrated solutions of the corresponding peptides in 5 mM Tris-Cl pH 7.5 with(out) 25 mM SDS were measured. The CD spectrometer (Jasco J-810) was operated at 25 °C using a quartz cuvette with a pathlength of 0.2 cm (Hellma Analytics 110-2-40). The spectra were recorded from 190-250 nm at scanning speed of 20 nm/min, band width of 1 nm and 5-fold data accumulation. After blank subtraction the mean data were converted to the molar ellipticity (deg*cm^2^/dmol) for each respective peptide. The percentage of peptide helicity was calculated with: helicity[%] = (−100*n*[Θ_222nm_])/(40,000*(n-4)), where n is the number of amide bonds in the peptide and Θ_222nm_ represents the mean molar ellipticity at 222 nm^38^.

### Fluorescence polarization-based binding experiments

All assays were performed at room temperature in low binding black 96-well microtiter plates (Greiner, 655 900) and measured as millipolarization (mP) units on a plate reader (Tecan Spark 20M). To determine the dissociation constant (*k*_D_), serial dilutions of GST-EloB/EloC-His protein (5.0 - 0 μM, 50 μL) in the assay buffer (25 mM Tris-Cl pH 7.5, 100 mM NaCl, 0.1 mM TCEP and 0.01% Triton X-100) were added to 100 μL of the fluorescently-tagged tracer peptide prepared in the same assay buffer. The final assay volume for each data point (triplicates) was 150 μL with 0.1 nM tracer. Each assay had two controls: blank (assay buffer alone) and 0.1 nM tracer in the assay buffer. The plates were incubated at room temperature to reach equilibrium and the mP values were measured after 60 min. The *k*_D_ values were calculated by converting the mP values into their corresponding anisotropy (A) values. These values were plotted versus the respective concentrations of the protein with a non-linear regression according to the following equation^39^:

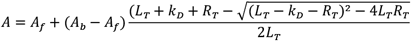

Hereby, *A* is the experimental anisotropy, *A*_*f*_ is the anisotropy for the free ligand, *A*_*b*_ is the anisotropy for the fully bound ligand, *L*_*T*_ is the total added concentration of ligand, *k*_D_ is the equilibrium dissociation constant and *R*_*T*_ is the total receptor concentration.

To determine the inhibitory constant (*k*_i_) competitive binding experiments were performed. GST-EloB/EloC-His protein was preincubated with tracer peptide in 25 mM Tris-Cl pH 7.5, 100 mM NaCl, 0.1 mM TCEP and 0.01% Triton X-100 at room temperature for 60 min to form the protein-tracer complex. To 142.5 μL of this protein-tracer complex, 7.5 μL of the corresponding peptide serial dilutions (120 – 0 μM) in DMSO were added, reaching a final concentration of 0.1 nM tracer, 1.9 nM protein and 5% DMSO. Three control wells were included in each experiment: blank (assay buffer alone), 100% inhibition (tracer alone) and 0% inhibition (protein-tracer complex), each with 5% DMSO. Inhibition constant (*k*_i_) values were calculated using the following equation described previously by Wang *et al*. and the standard deviation was calculated for each *k*_i_ value^40^.

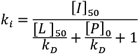

Hereby, *k*_i_ is the competitive inhibition constant, *[I]*_*50*_ is the concentration of free inhibitor at 50% inhibition, *[L]*_*50*_ is the concentration of free labelled ligand at 50% inhibition and *[P]*_*0*_ is the concentration of free protein at 0% inhibition.

### Hydrogen-deuterium exchange mass spectrometry (HDX-MS)

Protein stock solutions for HDX-MS contained GST-EloB/EloC complex (25 μM) either without (apo state) or with EPOP_wt_-R8, EPOP_mut_-R8 and EPOP_scr_-R8 peptide supplemented (50 μM final concentrations). Further preparation was aided by a two-arm robotic autosampler (LEAP Technologies) and conducted essentially as described previously with minor modifications^41^. In short, 7.5 μL of protein stock solution in the respective state was mixed with 67.5 μL D_2_O-containing buffer (25 mM Tris-Cl pH 7.5, 100 mM NaCl). After incubation at 25 °C for 10, 30, 100, 1,000 or 10,000 s, 55 μL were of the reaction was quenched with 400 mM KH_2_PO_4_/H_3_PO_4_, 2 M guanidine-HCl (pH 2.2) at 1 °C. From the resulting mixture 95 μL were injected into an ACQUITY UPLC M-Class System with HDX Technology (Waters)^42^. Non-deuterated samples were prepared similarly but in H_2_O-containing buffer. The injected samples were flushed out of the sample loop (50 μL) with constant flow (100 μL/min) of water + 0.1% (v/v) formic acid and guided to a cartridge (2 mm x 2 cm) filled with immobilized porcine pepsin, or a 1:1 mixture of immobilized protease type XVIII from *Rhizopus* sp. and protease type XIII from *Aspergillus saitoi* 12 °C. The resulting samples were combined and collected on an ACQUITY UPLC BEH C18 VanGuard Pre-column (1.7 μm, 2.1 mm x 5 mm (Waters)) at 0.5 °C. After 3 min, this column was placed in line with an ACQUITY UPLC BEH C18 column (1.7 μm, 1.0 × 100 mm (Waters)), and the peptides eluted at 0.5 °C with a gradient of water + 0.1% (v/v) formic acid (eluent A) and acetonitrile + 0.1% (v/v) formic acid (eluent B) at 60 μL/min flow rate as follows: 0-7 min/95-65% A, 7-8 min/65-15% A, 8-10 min/15% A. Eluting peptides were guided to a G2-Si HDMS mass spectrometer with ion mobility separation (Waters), and ionized by electrospray ionization (capillary temperature 250 °C, spray voltage 3.0 kV). Mass spectra were acquired over a range of 50 to 2,000 m/z in enhanced high definition MS (HDMS^E^) or high definition MS (HDMS) mode for non-deuterated and deuterated samples, respectively^43,44^. Three technical replicates (independent H/D exchange reactions) were measured per incubation time. Data were further analyzed as described^41^. Peptides were identified with ProteinLynx Global SERVER (PLGS, Waters) from the non-deuterated samples acquired with HDMS^E^. Deuterium incorporation into peptides was quantified with DynamX 3.0 software (Waters). Only peptides that were identified in the non-deuterated samples with a minimum intensity of 30,000 counts, a maximum length of 30 amino acids, a minimum number of three products with at least 0.1 product per amino acid, a maximum mass error of 25 ppm and a retention time tolerance of 0.5 minutes, were considered for analysis. Data obtained by HDX-MS are part of the Source Data supplied with this manuscript.

### Cell viability assays

Cells were grown in RPMI for PC3, DMEM for MCF7 and DMEM:F12 for SH-SY5Y, each with 10% fetal bovine serum (FBS) and 1% penicillin/streptomycin at 37 °C. During the assays the FBS content was lowered to 2.5%. Per well, 200 μL of PC3, MCF7 or SH-SY5Y at 10^5^ cell/mL were seeded into black 96-well cell culture microplates (Greiner, 655 086) and incubated 21 h. The next day 110 μL of the culture media was removed and 10 μL of peptides dissolved in ultrapure water (1.0 – 0 mM) were added. The plate was further incubated for 24 h. The cell viability for PC3 and MCF7 cells was assessed by resazurin fluorescence-based cell viability assay^45^: In each well, 20 μL of a 1.63 mM solution of resazurin in DPBS was added. The fluorescence at 590 nm was measured using a SpectraMax M5 plate reader in 30 min intervals until saturation in the untreated cells was reached. The background was subtracted and the slope of each individual well was calculated. Relative viability [%] = (slope_treated_)/(slope_untreated_) x 100.

The cell viability for SH-SY5Y cells was assessed by a luciferase-based cell viability assay: Per well, 100 μL of freshly-thawed and prepared CellTiter-Glo® reagent was added to each well and mixed according to the manufacturer’s instructions (CellTiter-Glo®, Promega) to induce cell lysis. The plate was incubated for 10 min and 150 μL of lysate was transferred to a white Lumitrac 96-well microplates (Greiner, 655 086). Luminescence was measured using the Tecan Spark 20M plate reader without filter and an integration time of 1,000 ms. The data were plotted and IC_50_ values were calculated by GraphPad Prism 6.

### Flow cytometry-based apoptosis detection

Into transparent sterile culture dishes, 12 mL of PC3 cells in RPMI supplemented with 2.5% FBS at a cell density of 60,000 cells/mL were seeded. The plates were incubated at 37 °C for 21 h. 100 μL of the culture media was removed and 100 μL of 300 μM peptides in ultrapure water were added resulting in a final concentration of 2.5 μM. Additionally, the dishes were treated with ultrapure water or 1 μM of doxorubicin as controls. After 24 h the addition was repeated for a total concentration of 5 μM peptide. After 24 h the culture medium was removed and the cells detached with 1 mL of accutase at room temperature for 15 min. The cells were transferred into a 1.5 mL centrifuge tube. From here on, all washing steps were performed by carefully resuspending the cells, centrifuging at 2,000 rpm for 2 min and removing the supernatant by liquid vacuum aspiration. The cells were washed with 500 μL DPBS and 500 μL of 1x Annexin V binding buffer and resuspended in 500 μL of 1x Annexin V binding buffer. Following the manufacturer’s protocol (BD Biosciences), 5 μL of Annexin V-FITC and 5 μL propidium iodide were added to the cell suspensions and incubated at room temperature for 15 min. The samples were analyzed with a BD LSRFortessa (Becton Dickinson Biosciences) flow cytometer within 1 h. For the excitation of the Annexin V-FITC and propidium iodide 488 nm and 561 nm lasers were used. The emission was detected using a 510/20 bandpass filter and a 632/22 bandpass filter. For each sample the data were collected until a total of 30,000 events were recorded. The obtained FCS data were analyzed using flowing software 2. Doublets exclusion was performed by gating in an SSC-W vs SSC-H plot and a subsequent FSC-W vs FSC-H plot. The final figures were prepared with a trial version of FlowJo™ v10.8 Software.

### Flow cytometry-based cell cycle analysis

PC3 cells were grown in RPMI medium supplemented with 1 % penicillin/streptomycin. Cells were seeded at a density of 10^5^ cells/mL and incubated with 2 mM Hydroxyurea (HU, Sigma-Aldrich) for 24h. HU was washed out thoroughly using PBS and control samples were collected, before lowering FCS concentrations to 2.5 % for PC3 peptide treatment with 6.5 μM of EPOP_wt_-R8, EPOP_mut_-R8, EPOP_scr_-R8 and R8 for 3h, 12h and 24h, respectively. After treatment, the culture medium was removed and the cells trypsinised at 37°C for up to 5 min. Detached cells were transferred to 15 mL tubes and washed twice with PBS. Afterwards, cells were re-suspended in 1 mL PBS and fixed drop wise under steady vortexing with 4 mL ice-cold 100 % ethanol. Samples were stored at -20°C until further processing. For propidium iodide (PI) staining, fixated cells were pelletized, re-suspended in PBS for up to 10 min, pelletized again, and finally re-suspended in 0.5 mL PI staining solution (50 μg/ml PI, 200 μg/ml RNase A, 0,1 % Triton-X in PBS). FACS analysis was performed with the BD Calibur (Becton Dickinson Biosciences) flow cytometer. For each sample, cell cycle data was collected until a total of 20,000 events were recorded. The obtained data was analyzed using flowing software 2. Histograms were prepared with a trial version of FlowJo™ v10.8 Software.

### Purification of GST-tagged Elongin B/C from bacteria for binding assays and HDX

For the expression of GST-ELOB and ELOC, *Escherichia coli* BL21(DE3) cells were transformed with the plasmid pRSF1Duet1 containing the coding sequences for GST-ELOB and ELOC. Cells were cultivated in baffled flasks at 37 °C and rigorous shaking until an optical density (at 595 nm) of 0.5 was reached, at which point protein overproduction was induced by addition of 0.5 mM IPTG. Cells were further grown overnight at 20 °C, harvested by centrifugation at 4,500 rpm (4 °C) and washed with PBS.

Cell pellets were resuspended in lysis buffer (1x HEMG [25 mM HEPES-K pH 7.6; 12.5 mM MgCl_2_; 5.1 mM EDTA; 10% glycerol], 500 mM NaCl, 0,1% NP-40; protease inhibitor cocktail (Roche) and 0.5 mM PMSF, as well as 1 mM DTT and 0.5 μg/mL lysozyme were added freshly). The mixture was rotated for 20 min head over tail at 4 °C. Then 1 μg/mL DNAse and RNAse were added and rotated for another 20 min. Cells were disrupted by sonication for 3x 30 sec. The extract was centrifuged for 15 min at 15,000 rpm. For further purification gluthation Sepharose 4B slurry (Cytiva) was washed three times with 1x HEMG supplemented with 500 mM NaCl and 0.1% NP-40, and the cell lysate containing GST-ELOB/C incubated with the resin for 2 h. Afterwards, beads were washed two times with lysis buffer containing 1 M NaCl (centrifugation at 3,000 rpm in between wash steps). In two further washing steps the buffer was exchanged to HEMG supplemented with 1 M NaCl and 0,01% NP-40. The protein was eluted with elution buffer (20 mM glutathione in HEMG supplemented with 1 M NaCl and 0.01% NP-40). The purified protein was dialyzed overnight in 25 mM Tris-Cl pH 7.5 using SnakeSkin^®^ialysis tubing (7 kDa molecular weight cutoff) and concentrated using Amicon™ Ultra-15 Centrifugal Filter Units (30 kDa molecular weight cutoff) (Merck) according to the requirements of the experiments.

### Pull-down experiment with bacterial GST-ELOC/B and FLAG-EPOP

For pull-down experiments GST-ELOC/B was expressed bacterially. FLAG-tagged EPOP was obtained from transfected HEK293 cells. For the expression of GST-ELOC/B the plasmid pRSF1Duet1-GST-ELOC-ELOB, which expresses GST-ELOC and untagged ELOB, was transformed into BL-21. A plasmid containing the GST (pGEX4T1) only was used as a control. Expression of proteins of interest was induced with 0.5 mM IPTG overnight at 20 °C. Bacteria were harvested by centrifugation at 4,000 rmp for 15 min at 4 °C and washed twice with cold PBS.

Pellets were mixed with Lysis buffer 1 [1x HEMG (25 mM HEPES-KOH pH 7.6; 12.5 mM MgCl_2_; 5.1 mM EDTA; 10% Glycerol), 200 mM NaCl, 0,1% NP-40; Protease inhibitor cocktail (Roche) and 0.5 mM PMSF, as well as 1 mM DTT and 0.5 μg/mL Lysozyme were added freshly]. The mixture was rotated for 20 min head over tail at 4 °C. Then 1 μg/mL DNAse and RNAse were added and the mixture was rotated for another 20 min. Cells were disrupted with sonication for 3x 30 sec. The extract was centrifuged at 15,000 rmp for 15 min. The supernatant containing ELOB/C was stored at -80 °C. GST-ELOB/C was purified using glutathione sepharose 4B slurry (Cytiva). Beads were prepared by washing three times with buffer 1 (1x HEMG with 200 mM NaCl, 0.1%. NP-40) The extract was added to the beads and rotated at 4 °C overnight. The next day beads bound with GST-ELOC/B were washed three times with buffer 2 (1x HEMG with 1 M NaCl, 0.1% NP-40).

To prepare the FLAG-EPOP extract from HEK293 cells, cells were seeded in 10 cm dishes with 2.2 × 10^7^ cells per dish. The next day, the expression vector pDEST-N-FLAG–EPOP was transfected using PEI. Two days later HEK293 cells were harvested. Cell pellet was dissolved in lysis buffer 2 (1x HEMG with 200 mM NaCl, 0.5% NP-40, 1x protease inhibitor cocktail (Roche), 0.5mM PMSF, 1 mM DTT).

For the pull-down beads bound with GST-ELOC/B were incubated for 1h with respective EPOP_wt_-R8, EPOP_mut_-R8, EPOP_scr_-R8 peptide (peptides were dissolved (1x HEMG with 200 mM NaCl, 0.1% NP-40) after 1 h extract containing FLAG-EPOP from HEK293 cells was added for another 1h. The final peptide concentration was 20 μM, 12.5 μM, or 6.5 μM.

Samples were washed three times with buffer 1. Samples were analysed via Western blot.

### Cell culture medium and handling

HEK293, HEK-T, PC3, MCF7, MIA PaCa-2, NCI-H23, and SH-SY5Y cells were grown at 37 °C with 5% (v/v) CO_2_. Growth media (RPMI for PC3 and NCI-H23, DMEM for, MIA PaCa-2 and MCF7, DMEM:F12/DMEM for SH-SY5Y, DMEM:F12 for HEK293 and HEK-T and MEM with 1% MEM NEAA for HEP G2) were supplemented with FBS (10% v/v), penicillin (100 units/mL), and streptomycin (100 μg/mL).

### Virus production and knockdown of ELOB and ELOC

To generate a knock-down of ELOB and ELOC in HEK-293, Hep G2, MCF7, MIA PaCa-2, NCI-H23, PC3, and SH-SY5Y, cells were subjected to lentiviral infection. A virus was produced in HEK-T cells by transfection of the packaging plasmids pMD2.g, psPAX2, and the respective Plko.1 vector, containing shRNA directed against control or ELOB and ELOC, respectively (shRNA Sequences in Supplementary Table S3) with PEI solution (1 mg/mL). The virus-containing supernatant was collected 48h after transfection. Cell lines were infected with virus for 48 h. Then cells were selected with 2 μg/mL of puromycin (SH-SY5Y (1 μg/mL)) for a further 48 h. Subsequently, cells were counted and seeded at a density of 6 × 10^5^ in a six-well plate and collected after 48 h for RNAseq of KD. Puromycin selection was stopped at this time point. For proliferation experiments, cells were seeded at 5 × 10^4^ per well.

### Proliferation assay

The cell viability was determined 1, 3, 5, and 7 days after seeding using an MTT assay (5 mg/mL Thiazolyl blue dissolved in PBS). Thereby, 90 μL reagent were added to the cell culture media in each well and incubated for 1-4 h. Stained cells were dissolved in 400 μL of lysis buffer (80% Isopropanol, 10% 1 M HCl, 10% Triton X-100). Absorption was measured at 595 nm using a plate reader (Microplate Reader, Molecular Devices, LLC, USA). All values were normalized to day 1 to compensate for variations in seeding density.

### RNA preparation and qPCR analysis

Cells were cultivated on 6-well plates up to 80-100% confluency for RNA isolation. RNA isolation was performed using the “RNeasy Mini Kit” (*Qiagen*) according to the manufacturer’s instructions. An additional on-column DNA digest (Qiagen) was added between the washing steps. According to the manufacturer’s manual, the Tetro cDNA Synthesis Kit was used for transcribing mRNA into cDNA. Subsequently, cDNA was diluted 1:20 with water to be used in RT-qPCR. For analysis by Real-Time quantitative PCR, the MyTaq™ Mix was used. For gene expression analysis, values were normalized to GAPDH expression (Primers in Supplementary Table S3). For RNA-seq of peptide-treated PC3 cells. Cells were seeded at 6 × 10^6^ into a six-well. 23 h later the medium was changed to RPMI containing only 2.5% FCS and the peptides EPOP_wt_-R8, EPOP_mut_-R8, and EPOP_scr_-R8 (dissolved in water), were added at a concentration of 12.5 μM. After 24 h cells were harvested.

For RNA-Seq, RNA integrity was assessed on Experion StdSens RNA chips (Bio-Rad). RNA-Seq libraries were prepared using the TruSeq Stranded mRNA Library Prep Kit (Illumina). RNA-Seq libraries were quantified on a Bioanalyzer (Agilent Technologies). Next-generation sequencing was performed on Illumina NextSeq550.

### Fractionation assay of peptide-treated cells

PC3 cells were treated for 24 h with 6.5 μM of the respective peptides and controls. During treatment concentration of FCS was lowered to 2.5%. Fractionation assays were conducted with the “Subcellular Protein Fractionation Kit for Cultured Cells” (Thermo Fisher) and performed as advised by the manufacturer. Protein concentration was estimated and equal amounts were loaded. ELOB/C as well as H3, Tubulin, and SP-1^46^ were detected via Western Blotting (see Supplementary Table S2 for antibodies).

### Immunofluorescence staining

For immunofluorescence staining, HEK293 cells were seeded on coverslips. Cells were fixed with 4% formaldehyde (*w*/*v*), methanol-free (Thermo Fisher Scientific, Waltham, MA, USA; PI28906), and subsequently permeabilized with 0.5% Triton X-100 in PBS. Blocking was performed with 10% FBS in PBS. To detect ELOB and ELOC, the respective primary antibodies (Supplementary Table S2) were diluted at 1:500 in the blocking solution. After primary antibody incubation, the cells were washed three times with 0.5% Triton X-100 in PBS. Secondary antibody incubation was conducted using Alexa Fluor 488 and goat anti-rabbit IgG (H+L) (Thermo Fisher Scientific, Waltham, MA, USA; A-11008) at a 1:1,000 dilution. Following three washing steps, the coverslips were mounted onto microscopy slides using VECTASHIELD^®^ Antifade Mounting Medium with DAPI (VECTOR Laboratories).

### Colony formation assay

To examine the ability of cells to form colonies, the cells were seeded at low density (1 × 10^3^ cells per well on 6-well plates) and cultured for 10 days. Thereby cells were treated with 6.5 μM of the respective peptides. During the treatment, the FCS content of the medium was lowered to 2.5%. After 24 h, the medium was replaced by one containing 10% FCS. For analysis, cells were washed once with PBS and then fixed with 100% methanol for 20 min. Afterward, the cells were stained for 5 min with 0.5% crystal violet in 25% methanol. To remove excess color, the plates were washed with de-ionized water until single colonies were visible. Images were taken, and colonies were counted using ImageJ Fiji (v1.53p). The mean value of three biological replicates was determined.

### Mass spectrometry

To identify potential EPOP interaction partners via mass spectrometric analysis, nuclear extracts were prepared for a FLAG-IP. Per co-immuniprecipitation experiment 1-2 × 10^8^ HEK-293 cells stably expressing either FLAG-HA-EPOP(WT); FLAG-HA-EPOP(L40A), FLAG-HA-EPOP(ΔC-ter.) and FLAG-HA-GFP as control were used. After collection, cells were centrifuged at 2,000 rpm and 4 °C for 10 min. Cells were resuspended in 5-times packed cell volume(PV) hypotonic buffer (10 mM Tris-Cl pH 7.3, 10 mM KCl, 1.5 mM MgCl_2_ 0.2 mM PMSF, 10 mM ß-mercaptoethanol 1 protease inhibitor cocktail tablet (Roche) and shaken in the thermomixer at 4 °C for 10-15 min. The now-swollen cells were resuspended in 5 mL hypotonic buffer and transferred to a dounce homogenizer where cells were disrupted to obtain the nuclei. To remove cell debris, lysates were centrifuged at 3,500 rpm and 4 °C for 15 min. The pellet was resuspended in 1x pellet volume low alt uffer (20 mM Tris-Cl pH 7.3, 20 mM KCl, 1.5 mM MgCl_2,_ 0.2 mM EDTA, 25% glycerol, 0.2 mM PMSF, 10 mM ß-mercaptoethanol, 1x protease inhibitor cocktail). Cells were dounced for 7-10x, and then 0.66x pellet volume of high salt buffer (20 mM Tris-Cl pH 7.3, 1.2 M KCl, 1.5 mM MgCl_2_, 0.2 mM EDTA, 25% glycerol, 0.2 mM PMSF, 10 mM β-mercaptoethanol) was added gradually, while constantly mixing. The lysate was incubated whilst rocking for 30 min at 4 °C. It was then centrifuged at 13,300 rpm and 4 °C for 15 min and the supernatant was transferred to a “Slide-A-Lyzer™ G2 Dialysis Cassettes” (3.5K) (Thermo Scientific) and dialyzed against 3 L of dialysis buffer (20 mM Tris-Cl pH 7.3, 100 mM KCl, 0.2 mM EDTA, 20% glycerol, 0.2 mM PMSF, 1 mM DTT) overnight.

To perform the FLAG-IP, the material was retrieved from the dialysis chambers and centrifuged at 13,300 rpm and 4 °C for 30 min. The ANTI-FLAG M2 affinity gel (Sigma) was prepared by washing once with TAP buffer (50 mM Tris-Cl pH 7.9, 100 mM KCl, 5 mM MgCl_2_, 0.2 mM EDTA, 10% glycerol, 0.2 mM PMSF, 1mM DTT), three times with 100 mM glycine (pH 2.5), once with 1 M Tris-Cl (pH 7.9) and finally once with TAP buffer. The extract was added to the beads and rotated at 4 °C for 4 h. Afterwards, beads were washed three times with TAP buffer to wash away unbound material.

From there on, two different protocols were employed depending on whether the material was to be analyzed by mass spectrometry or on a silver-stained gel. In the former case, the beads were washed three times in 50 mM ammonium bicarbonate (NH_4_HCO_3_). The IP-ed material was then sent in for mass spectrometry analysis at the Biomedical Center Munich, protein analysis unit (Head: Axel Imhof).

If the material was to be loaded on a silver gel or used for western blotting, the beads were incubated with 0.2 mg/mL single FLAG-peptide(Sigma) in TAP buffer and rotated at 4 °C for 1 h. Afterwards, the beads were centrifuged at 3,000 rpm at 4 °C for 5 min for elution. The supernatant, containing the immunoprecipitated material, was then used for downstream applications.

### Flag-IP after peptide treatment

All ectopic co-immunoprecipitation (Co-IP) experiments were performed in HEK293 cells. Cells were seeded in 10-cm dishes with 2.2 × 10^7^ cells per dish. After 24 h, the expression constructs encoding for N-FLAG–tagged proteins were transfected using PEI. For Co-IP with FLAG-EPOP, a stable line was made by selecting the cells with 2 μg/mL puromycin. FLAG-VHL was transfected transiently.

The next day, cells were washed with PBS and the medium changed to one containing only 2.5% FCS. The peptides EPOP_wt_-R8, EPOP_mut_-R8, and EPOP_scr_-R8 were dissolved in water and cells treated for 24 h with the concentrations 12.5, 6.25, and 3.125 μM of the peptide.

Two days after transfection, cell lysis was performed using Co-IP lysis buffer (50 mM Tris-Cl pH 7.5, 150 mM NaCl, 1% Triton X-100, 1 mM EDTA, 10% glycerol, 1x protease inhibitor cocktail EDTA-free (Roche), and 0.5 mM PMSF). Cells were shaken for 30 min at 4 °C followed by centrifugation for 10 min at 13,000 rpm at 4 °C. The extract was immediately used for FLAG-IP. Beads were equilibrated by washing three times with Co-IP lysis buffer. To bind FLAG-tagged proteins, extracts were incubated with 40 μL equilibrated ANTI-FLAG M2 Affinity Gel (Merck, A2220) for approximately 3 hours at 4 °C. After incubation, three washing steps with Co-IP wash buffer were performed (50 mM Tris-Cl pH 7.4, 250 mM NaCl, 1 mM EDTA, 1% Triton X-100, 10% glycerol). The FLAG beads were prepared for Western blotting through the addition of 2x Laemmli buffer. Detection of proteins in the input, supernatant, and IP fractions was conducted via Western blotting. Co-IP experiments were repeated at least three times.

### Bioinformatic analysis

RNA-Seq samples were aligned to the human transcriptome GENCODE v32 using RNA-Star (2.7.8a)^47^. Reads per gene were calculated using feature counts (2.0.1). Differentially regulated genes and normalized read counts were determined using DeSeq2 (2.11.40.7)^48^. PCA Analysis was performed using Bioconductor/R using the DeSeq2 package^49^. Genes with at least 0.75-fold (Knockdown) or 0.35-fold (peptide treatment) (log2) difference and a *p*-value below 0.01 were considered differentially expressed genes. Gene set enrichment analysis was performed using standard settings^21^ and visualised via ggplot.

### Statistics

All in vitro and cell-based experiments were performed in three independent experiments using new reagents and/or cells from another passage. Each data point represents a technical triplicate. For flow cytometry experiments three independent measurements were conducted for subsequent statistical analysis. All calculated values are given as the mean ± standard deviation (s.d.) or standard error (s.e.). Dose-response curves were fit to four parameter logistic (4-PL) curve using GraphPad 6 (log[inhibitor] vs response – variable slope; equation: Y=Bottom+(Top-Bottom)/(1+10^((LogIC50-X)×HillSlope)^). For comparison of multiple groups with one variable, one-way ANOVA variance analysis was performed. Statistical analyses were conducted using GraphPad Prism 6.0. Briefly, pairwise comparisons were performed by using two-tailed unpaired t-tests. Specific statistical tests are identified in the corresponding figure legends. *p* < 0.05 was considered significant.

## Data availability

The RNA-Seq data have been deposited to the GEO repository under the accession number GSEXXXXXXX. The proteomic data were uploaded to the PRIDE Repository under the Accession number XXXXX. All other data supporting the findings of this study are available within the article and its supplementary information files and from the corresponding author upon reasonable request.

## Acknowledgements

We thank Prof. Dr. Zhanxin Wang for providing expression constructs, Prof. Dr. Uta-Maria Bauer and Prof. Dr. Eric. Meggers for the generous use of her facilities. We thank Prof. Dr. Michael Wanzel for supplying MCF7 cells and Dr. Silvia Gonzalez Sierra for flow cytometry acquisition, interpretation and support. We thank BD Bioscience for the FlowJo software access. We thank the cellular imaging core facility at the University of Marburg for their advice and use of facilities. This work was supported by following the DFG programs: Investigator program GO1011/11-1 (425970020) to O.V., TRR81: ‘Chromatin Changes in Differentiation and Malignancies’ TRR81/3, A17, Z04 (109546710) to R.L. and O.V, core facility for interaction, dynamics and assembly of biomolecular structures (324652314) to G.B., and AB 792/1-1 (423428279) for F.A.

## Author information

### Authors and Affiliations

#### Contributions

V.T.T. synthesised the peptides, purified ELOB/C, conducted the FP-, cell-based assays as well as flow cytometry and contributed to HDX interpretation; F.A. contributed to ELOB/C purification and cell-based assays. W.S. performed HDX analysis and interpretation; G.B. performed HDX interpretation; S.F. purified ELOB/C, and performed cellular and molecular experiments. C.S. and L.M.W. contributed to cellular and molecular experiments. R.L. performed the bioinformatic analysis. I.F. performed semi-quantitative mass spectrometry analysis. A.N and T.S performed and supervised the next generation sequencing experiments. O.V. and R.L conceived of the idea for the study, designed, directed and interpreted experiments, co-supervised the work and fulfilled the role of corresponding authors. V.T.T, S.F., W.S., R.L. and O.V. wrote the manuscript. All authors commented on and approved the manuscript.

### Corresponding author

Correspondence to Robert Liefke or Olalla Vázquez.

## Extended Figures

**Extended Data Fig. 1.**
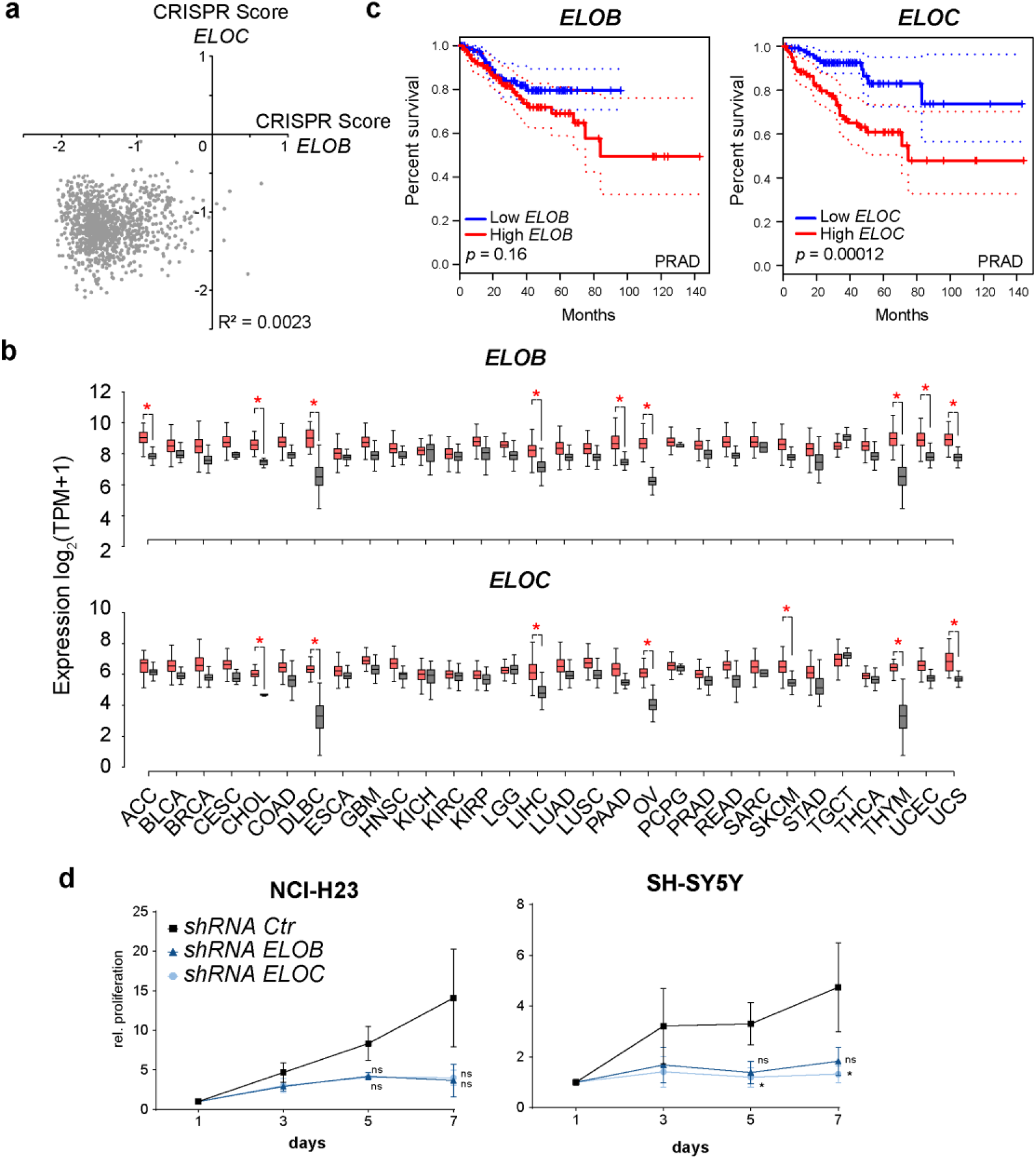
Role of ELOB/C in cancer. **a**, Comparison of CRISPR Scores for *ELOB* and *ELOC* from individual cancer cell lines^17,18^. **b**, Comparison of *ELOB* and *ELOC* expression in normal and cancer tissue. Data from TCGA and GEPIA^19,20^. The whisker-box plots represent the lower quartile, median and upper quartile of the data with 5% and 95% whiskers. Asterisks indicate significant changes *p* < 0.05. **c**, Kaplan-Meier Survival Plot (Disease free survival) in Prostate cancer adenocarcinoma (PRAD) based on expression of *ELOB* or *ELOC*. Data from TCGA^19^ and visualised by GEPIA^20^. **d**, Proliferation of cancer cell lines NCI-H23 and SH-SYS5 after *ELOB* or *ELOC* knockdown. Three independent replicates were collected and a ratio paired t-test of paired samples was conducted at the 5^th^ and 7^th^ day. Gaussian distribution was assumed as well as consistent ratios of paired values (n.s., *p* > 0.05; *, *p* < 0.05; **, *p* < 0.01; ***, *p* < 0.001).

**Extended Data Fig. 2.**
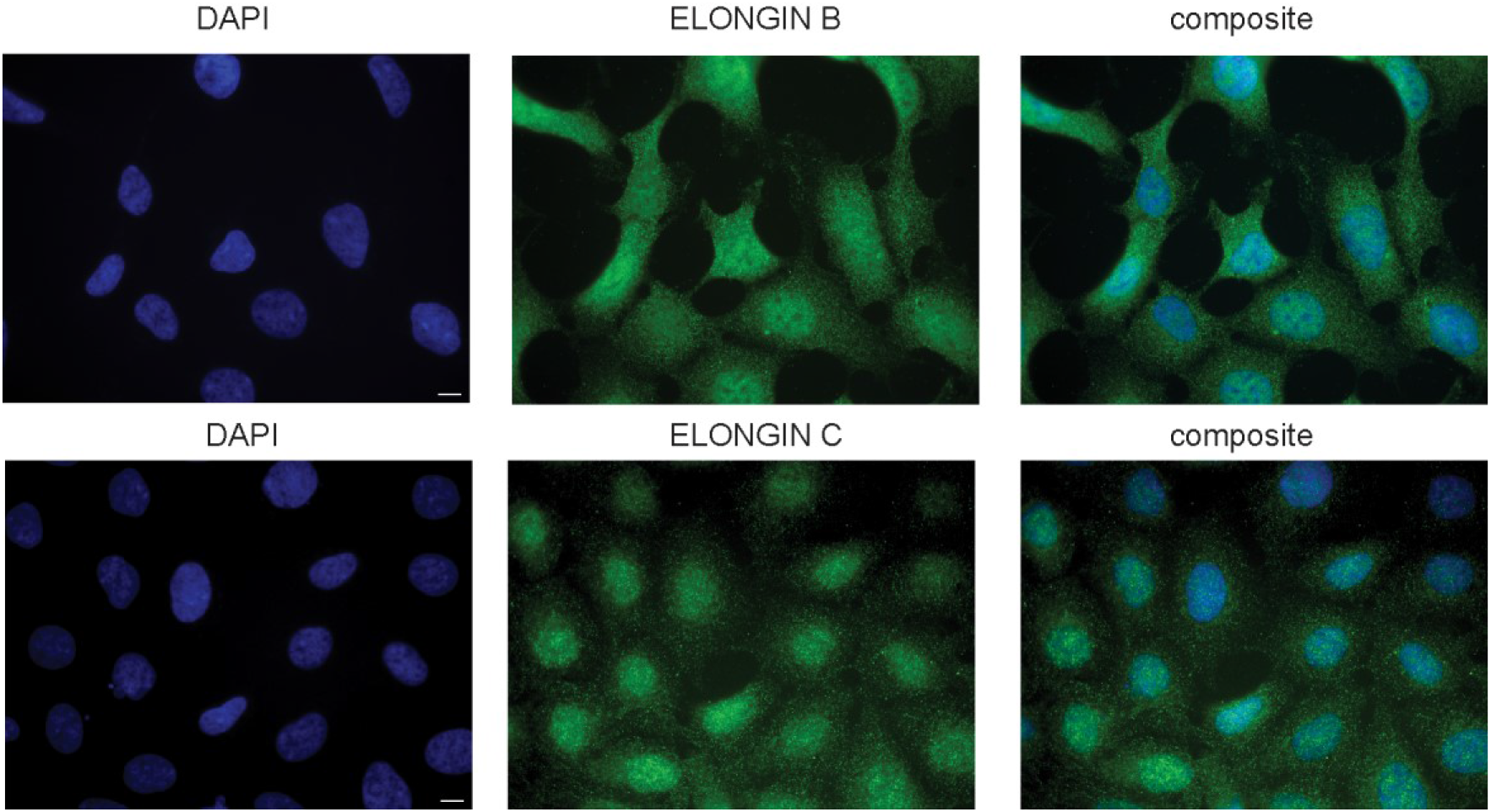
Immunofluorescence of endogenous ELOB and ELOC in HEK293 cells. Analysis of nuclear localisation of ELOB and ELOC using specific antibodies.

**Extended Data Fig. 3.**
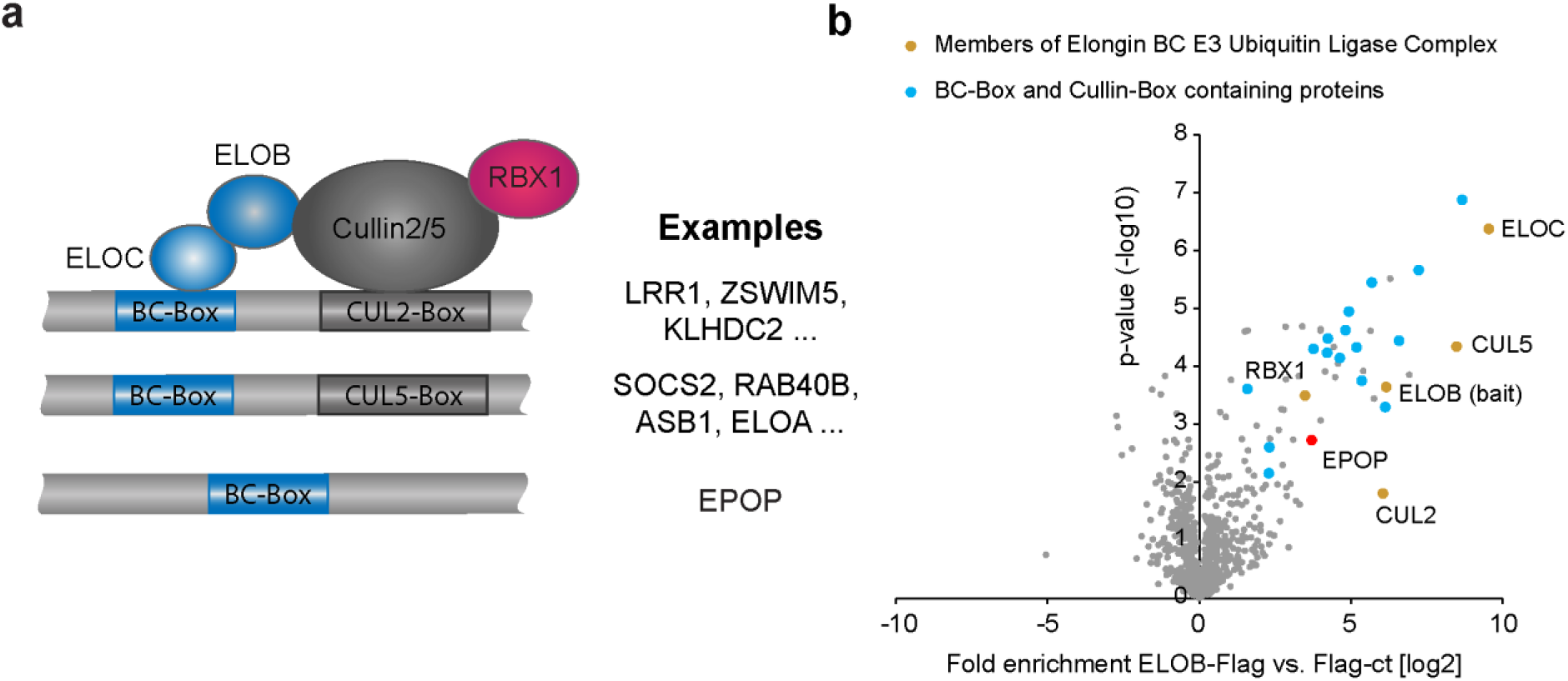
EPOP is a unique ELOB/C interacting protein. **a**, Overview showing the motif composition of typical BC-Box containing proteins. The names of some example proteins are presented. **b**, Re-analysis of previously published mass-spectrometry data analysing the interactome of ELOB^15^. Yellow colour marks core components of the ELOB/C-associated E3 ubiquitin ligase complex. Blue color denotes proteins that possess both a BC-box and a cullin-box. EPOP is shown in red.

**Extended Data Fig. 4.**
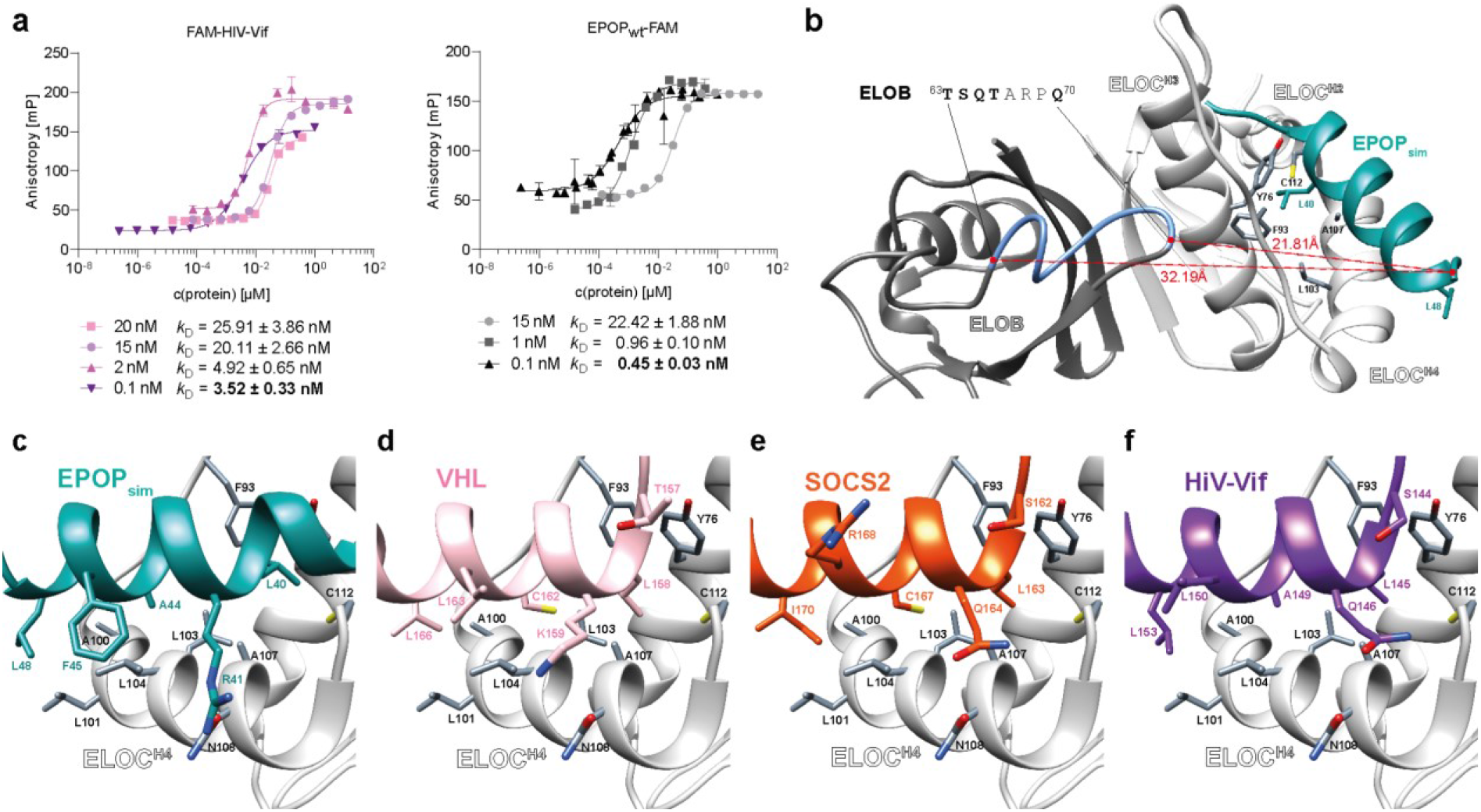
Tracer-dependent *k*_D_ overestimation of tight binders FAM-HIV-Vif and EPOP_wt_-FAM. Structural analysis and comparison of BC-boxes. **a**, Fluorescence polarization-based binding experiments of EPOP_wt_-FAM and FAM-HIV-Vif at varying concentrations (20 - 0.1 nM) to GST-ELOB/ELOC in 25 mM Tris pH 7.5, 100 mM NaCl, 0.1 mM TCEP and 0.02% Triton X-100. **b**, Unspecific interaction caused by ionic R8-linker potentially affects H/D-exchange of ELOB. Approximate distance of the linker to polar sequence ELOB^63-70^ (blue) indicated with dotted line (red). **c**, Close-up of ELOC^H4^ with predicted EPOP_sim_. **d**, Close-up of ELOC^H4^-pVHL interaction surface (1LM8)^25^. **e**, Close-up of ELOC^H4^-SOCS2 interaction surface (2C9W)^26^. **f**, Close-up of ELOC^H4^-HIV-Vif interaction surface (3DCG)^23^.

**Extended Data Fig. 5.**
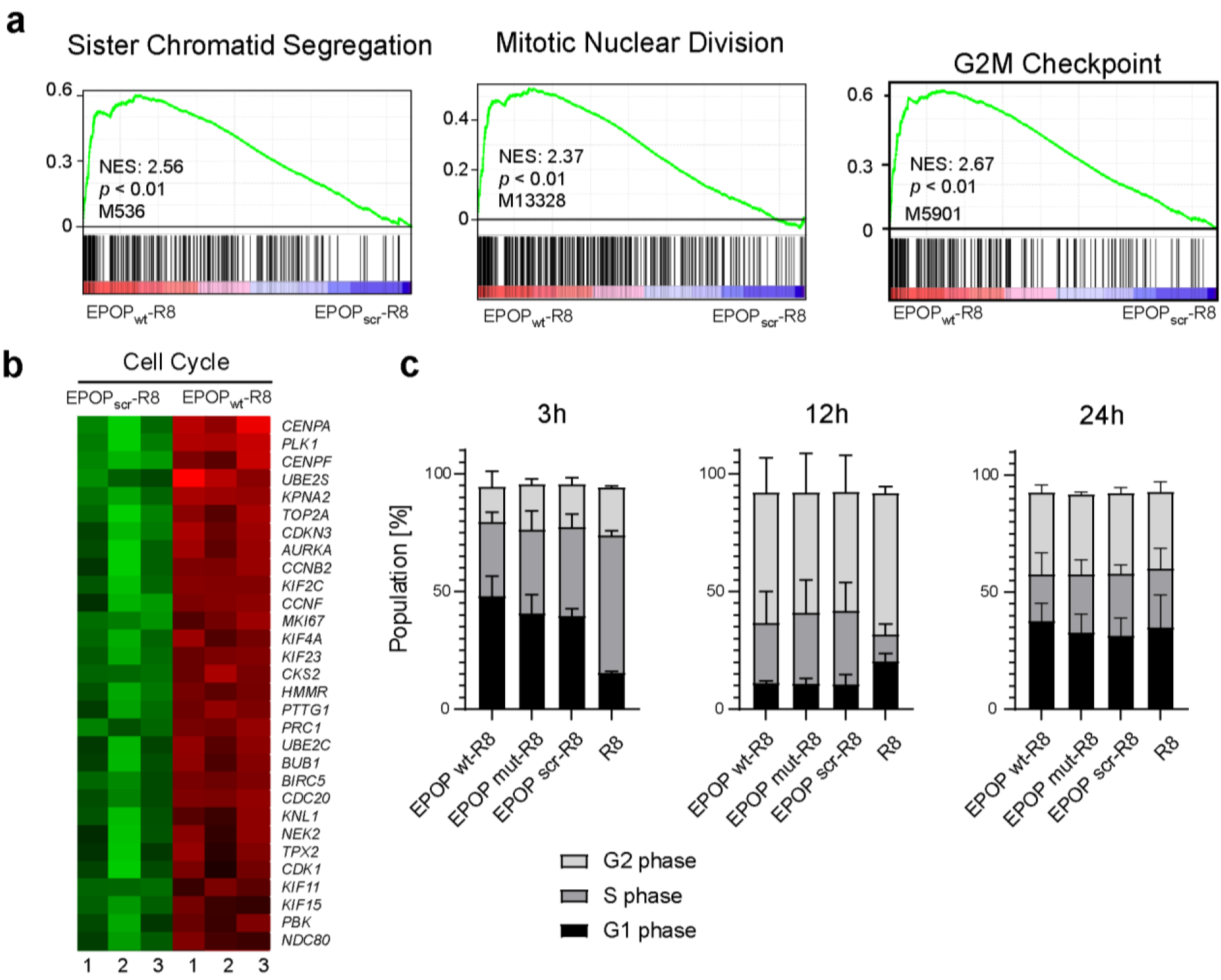
Peptide treatment influences cell cycle genes in PC3 cells. **a**, GSEA analysis showing the upregulation of cell cycle-related pathways upon peptide treatment. **b**, Heatmap showing the upregulation of key cell cycle genes upon peptide treatment. **c**, Cell cycle analysis of PC3 cells treated with 6.5 μM EPOP BC-box peptides over 3h, 12h and 24h, respectively, determined by propidium iodide staining in flow cytometry experiments. Data represent mean ± s.d. of n = 3 experimental independent replicates. Respective histograms are presented in Supplementary Fig. S22.

**Extended Data Fig. 6.**
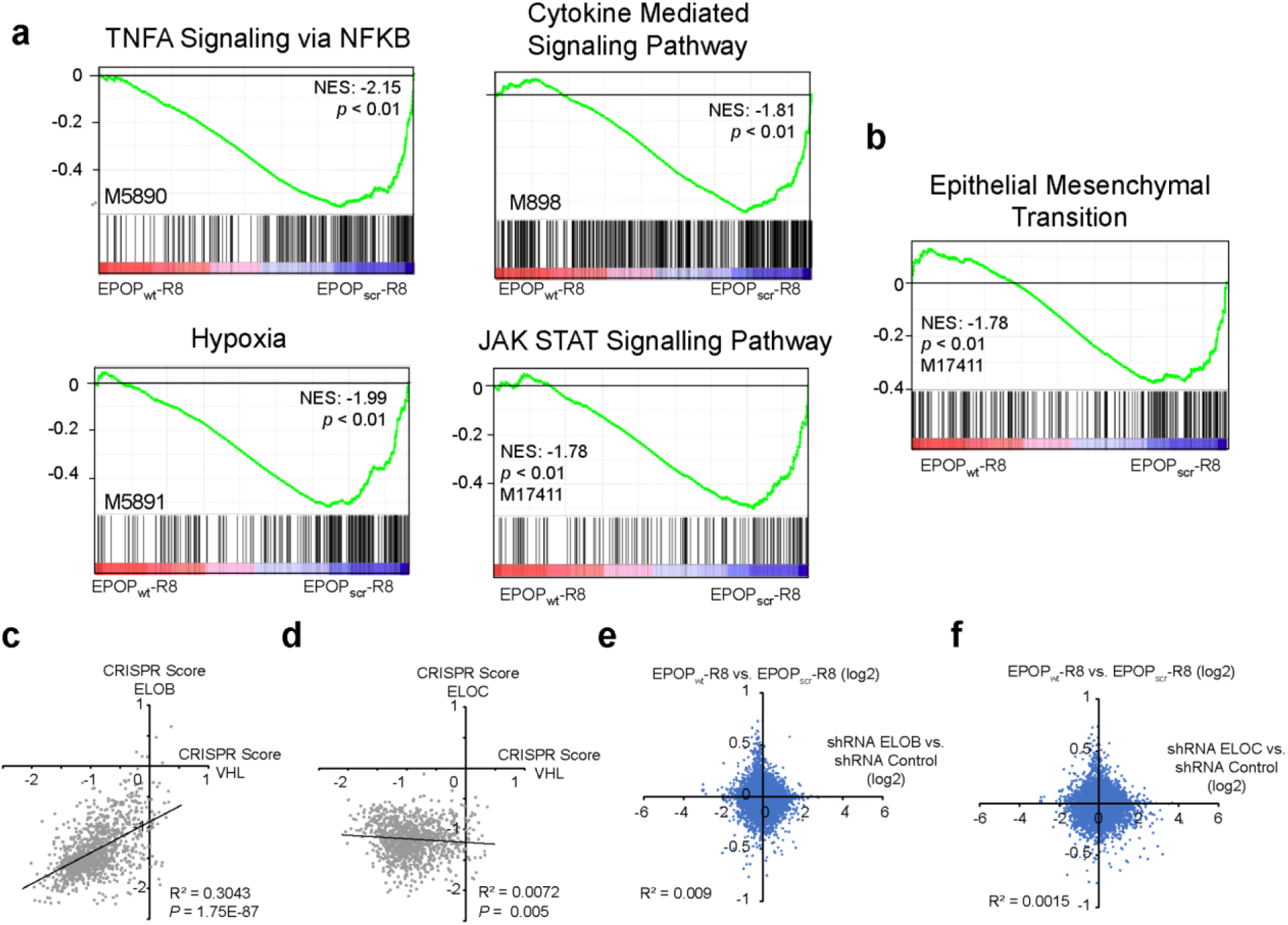
GSEA analysis upon peptide treatment and comparison to knockdown. **a**, GSEA analysis showing the downregulation of several signalling pathways. **b**, GSEA shows the downregulation of EMT-related genes upon peptide treatment. **c-d**, Correlation of CRISPR scores from *VHL* with *ELOB* (c) and *ELOC* (d) in cancer cell lines, downloaded from GEPIA^20^. **e-f**, Correlation of gene expression changes upon peptides treatment (Fig. 6f) versus knockdown (Fig. 1b) of *ELOB* (e) or *ELOC* (f).

