## Supplementary Information for "Peptide-mediated inhibition of the transcriptional regulator Elongin BC induces apoptosis in cancer cells"

2.1 Solid-Phase Peptide Synthesis 6

2.2 Hydrogen-Deuterium Exchange Mass Spectrometry (HDX-MS) 11

2.3 Cell Viability Assays 13

2.4 Replicates of apoptosis assays 14

2.5 Silver for mass spectrometry 15

2.6 Replicates of Co-Immunoprecipitation experiments 16

2.7 Histograms of cell cycle analysis 16

2.8 Antibodies and Oligonucleotides 17

#### Abbreviations

| | |
|---|---|
| Aba | 4-acetamidobenzoic acid |
| $Ac_2O$ | acetic anhydride |
| Ahx | 6-aminohexanoic acid |
| Alloc | allyloxycarbonyl |
| Boc | *tert*-butylcarbonyl |
| calcd. | calculated |
| Da | dalton |
| DBPS | Dulbecco's phosphate buffered saline |
| DIC | *N*,*N'*-diisopropylcarbodiimide |
| DIPEA | *N,N*-diisopropylethylamine |
| DMEM | Dulbecco's Modified Eagle Medium |
| DMEM:F12 | Dulbecco's Modified Eagle Medium Mixture F-12 |
| DMF | dimethylformamide |
| DMSO | dimethyl sulfoxide |
| DOX | doxorubicin |
| DTT | dithiothreitol |
| EDT | ethane-1,2-dithiol |
| ESI | electrospray ionization |
| FAM | 5(6)-carboxyfluorescein |
| FBS | fetal bovine serum |
| Fmoc | 9-fluorenylmethoxycarbonyl |
| FP | fluorescence polarization |
| $H_3PO_4$ | phosphoric acid |
| HATU | 1-[bis(dimethylamin)methylen]-1H-1,2,3-triazol[4,5-b]pyridinium 3-oxidhexafluorphosphat |
| HCl | hydrochloric acid |
| HPLC | high-performance liquid chromatography |
| HRMS | high resolution mass spectrometry |
| $IC_{50}$ | half inhibitory concentration |
| $k_d$ | equilibrium dissociation constant |
| $KH_2PO_4$ | potassium dihydrogen phosphate |
| MeCN | acetonitrile |
| mP | milli polarization value |
| mAU | milli arbitrary unit |
| m/z | mass-to-charge ratio |
| NMM | *N*-methylmorpholin |
| NMP | *N*-methyl-2-pyrrolidon |
| PBS | phosphate buffered saline |
| RFU | relative fluorescence unit |

| | |
|---|---|
| RP-HPLC | reversed phase high-performance liquid chromatography |
| RPMI | Roswell Park Memorial Institute 1640 Medium |
| rt | room temperature |
| SDS | sodium dodecyl sulfate |
| TCEP | tris(2-carboxyethyl)phosphine |
| TFA | trifluoroacetic acid |
| $t_R$ | retention time |
| Tris | tris(hydroxymethyl)aminomethane |
| UV-vis | ultraviolet–visible |
| λ | wavelength |

### 1 Material

All commercially available reagents were purchased from the following companies, and used without further purification: *N*-methylmorpholine (NMM), thioanisol, 1,2-ethandithiol (EDT), Triton X-100, trypan blue, trypsin-EDTA, penicillin-streptomycin biograde dimethylsulfoxide (DMSO), ANTI-FLAG® M2-Affinitätsgel (A2220), FLAG® Peptid (F3290) and accutase solution (A6964) from Sigma Aldrich (USA); ≥ 99.9% sodium chloride (NaCl) , piperidine, tris(2-carboxyethyl)phosphine (TCEP), tris(hydroxymethyl)aminomethane (Tris) ,2 mL polypropylene reactors with plunger and frit pore size 25 µm (7926.1), *N,N*-diisopropylethylamine (DIPEA) and ROTI®Black P for silver staining from Carl Roth (Germany); 5(6)-carboxyfluorescein (FAM), 4-acetamidobenzoic acid (Aba), 2,6-lutidine, palladium acetate, phenylsilane, trifluoracetic acid (TFA) and glycerol from Acros (USA); frits 0.5”/0.118” frits OD/THK with 25-40 µm pores 26030301 for Apex 396 from Aapptec, LCC (USA); microscale columns 35.091 from Intavis (Germany); Fmoc-protected amino acids, peptide grade dimethylformamide (DMF), TentaGel S RAM resin, Fmoc-Gly-OH, Fmoc-L-Asn(Trt)-OH, Fmoc-L-His(Trt)-OH, Fmoc-L-Ile-OH, Fmoc-L-Lys(Alloc)-OH, Fmoc-L-Met-OH, Fmoc-L-Phe-OH, Fmoc-L-Pro-OH*$H_20$, Fmoc-L-Thr(tBu)-OH, Fmoc-L-Tyr(tBu)-OH, Fmoc-L-Val-OH and 1-[bis(dimethylamino)methylene]-1H-1,2,3-triazolo[4,5-b]pyridinium-3-oxid hexafluorophoshate (HATU) from Iris Biotech (Germany); Fmoc-L-Ala-OH-OH, Fmoc-L-Arg(Pbf)-OH, Fmoc-L-Cys(Trt)-OH, Fmoc-L-Gln(Trt)-OH, Fmoc-L-Glu(OtBu)-OH, Fmoc-L-Leu-OH, Fmoc-L-Lys(Boc)-OH, Fmoc-L-Ser(tBu)-OH, *N,N′*-diisopropylcarbodiimid (DIC), and OxymaPure from Carbolution (Germany); Fmoc-e-Ahx-OH, HPLC grade acetonitrile, molecular biology grade mono- and dibasic potassium phosphate from Merck (Germany); Tetro Reverse Transcriptase (BIO-65050) from Meridian Bioscience; fetal bovine serum (FBS) from Capricorn Scientific (USA); resazurin sodium salt, Roswell Park Memorial Institute Media 1640 (RPMI), Dulbecco's Modified Eagle Medium (DMEM), Gibco Dulbecco's Modified Eagle Medium Mixture F-12 (DMEM:F12), Dulbecco′s Phosphate Buffered Saline (DPBS) and Subcellular Protein Fractionation Kit for Cultured Cells from ThermoFisher; Opti-MEM® I (1X) + GlutaMAXTM -I; MEM (1x) and MEM NEAA (100X) from Gibco;sterile culture dishes with ventilation cams (92 x16 mm), T75 flat-bottomed culture flasks from Sarstedt (Germany); 96 well cell culture microplates black with lid (655 086), 96 well microplates F-bottom white Lumitrac (655 075) and F-bottom black non-binding (655 900) from Greiner Bio-One (Austria); CellTiter-Glo® luminescent cell viability assay (G7570) from Promega (USA); Annexin V and propidium iodide apoptosis detection kit from BD Biosciences (Germarny); RNeasy Mini Kit (74104) as well as DNase, RNase free (79254) were purchased from Quiagen; Ultrapure water of type 1 was obtained with a MicroPure Water Purification System from TKA (Germany).

### 2 Supplementary Figures and Tables

#### 2.1 Solid-Phase Peptide Synthesis

**Supplementary Table S1:** Columns for analytical and (semi-)preparative HPLC-MS. Column 1 and column 2 were used for the semi-preparative or preparative purification of peptide crude while column 3 was for the characterization of the final purified peptides.

| Column | Type | Dimensions | Flow | Purpose |
|---|---|---|---|---|
| Column 1 | Jupiter C18 300 Å column (Phenomenex) | 250 x 10 mm, 10 µm | 6 mL/min | semi-preparative |
| Column 2 | Nucleodur EC 125/2 100-C18 (Macherey & Nagel) | 250 x 21 mm, 5 µm | 10 mL/min | preparative |
| Column 3 | Eclipse XBD-C18 (Agilent Technologies) | 150 x 4.6 mm, 5 µm | 1 mL/min | analytic |

**EPOP$_{wt}$-FAM**

$H_2N$-EFSPLCLRALAFCALAK-Ahx-Ahx-K(FAM)-$CONH_2$; peptide was synthesized in 5 µmol scale. After purification (column 1, 5-75% MeCN) , the 3 x TFA salt product (5.95 mg, 2.04 µmol, 41% yield) was obtained as a white solid. $t_R$ = 15.39 min. Purity ≥ 97%. Formula: $C_{124}H_{182}N_{26}O_{29}S_2$. Molecular weight: 2565.09 HRMS-ESI+ (m/z): $[M+3H]^{3+}$ calcd.: 855.7752; found: 855.7742.

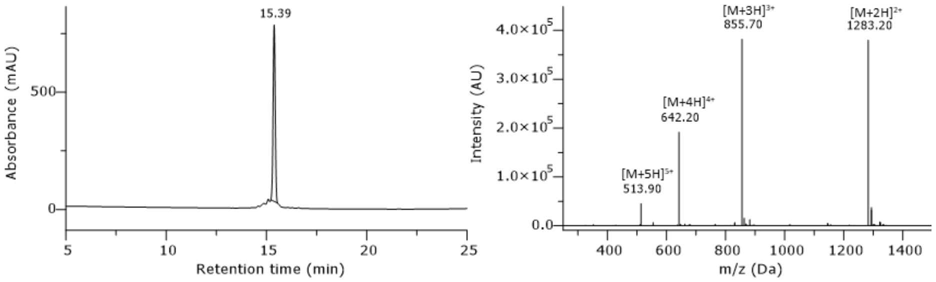

**Supplementary Fig. S1:** HPLC chromatogram of purified peptide EPOP$_{wt}$-FAM. Gradient 5-95% B in column 3, monitored at 220 nm.

**FAM-HIV-Vif**

FAM-HNKVGSLQYLALTALI-$CONH_2$; peptide was synthesized in 5 µmol scale. After purification (column 1, 5-75% MeCN), the 2 x TFA salt product (3.2 mg, 1.38 µmol, 28% yield) was obtained as white solid. $t_R$ = 18.22 min. Purity 95%. Formula: $C_{101}H_{144}N_{22}O_{27}$. Molecular weight: 2098.39. HRMS-ESI+ (m/z): $[M+2H]^{2+}$ calcd.: 1050.0373; found: 1050.0380.

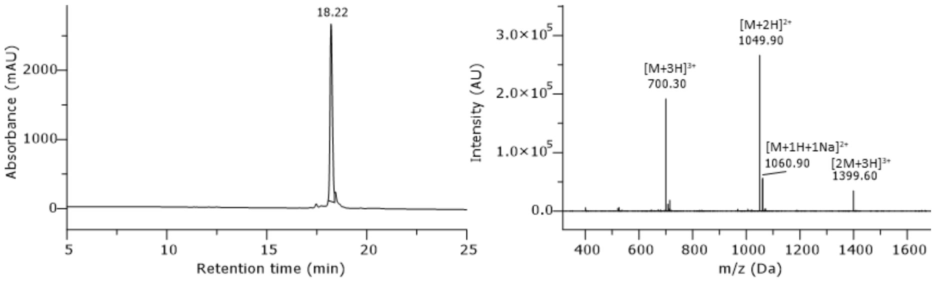

**Supplementary Fig. S2:** HPLC chromatogram of purified peptide FAM-HIV-Vif. Gradient 5-95% B in column 3, monitored at 220 nm.

**EPOP$_{wt}$**

$H_2N$-EFSPLCLRALAFCALAK-Ahx-Ahx-K(Aba)-$CONH_2$; peptide was synthesized in 5 µmol scale. After purification (column 1, 5-75% MeCN), the 3x TFA salt product (5.1 mg, 1.88 µmol, 38% yield) was obtained as white solid. $t_R$ = 18.17 min. Purity ≥ 97%. Formula: $C_{112}H_{179}N_{27}O_{25}S_2$. Molecular weight: 2367.95. HRMS-ESI+ (m/z): $[M+3H]^{3+}$ calcd.: 790.1085; found: 790.1095.

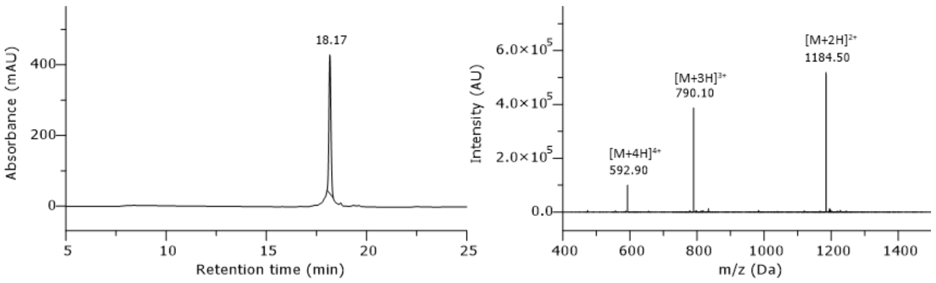

**Supplementary Fig. S3:** HPLC chromatogram of purified peptide EPOP$_{wt}$. Gradient 5-95% B in column 3, monitored at 220 nm.

**EPOP$_{mut}$**

$H_2N$-EFSPLC**A**RALAFCALAK-Ahx-Ahx-K(Aba)-$CONH_2$; peptide was synthesized in 5 µmol scale. After purification (column 1, 5-75% MeCN), the 3 x TFA salt product (3.6 mg, 1.35 µmol, 27% yield) was obtained as white solid. $t_R$ = 17.25 min. Purity ≥ 99%. Formula: $C_{109}H_{173}N_{27}O_{25}S_2$. Molecular weight: 2325.87. HRMS-ESI+ (m/z): $[M+3H]^{3+}$ calcd.: 776.0928; found: 776.0917.

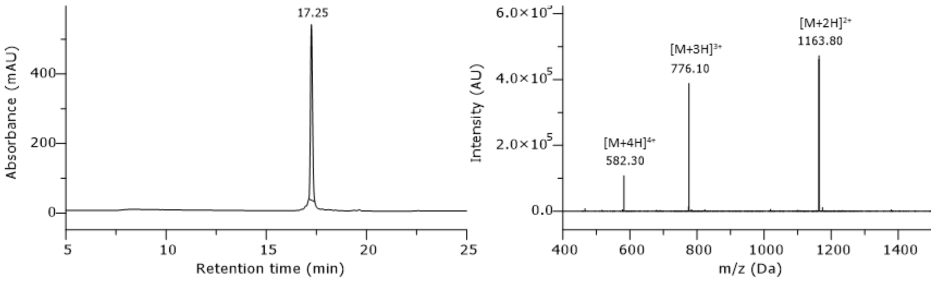

**Supplementary Fig. S4:** HPLC chromatogram of purified peptide EPOP$_{mut}$. Gradient 5-95% B in column 3, monitored at 220 nm.

**EPOP$_{scr}$**

$H_2N$-CLARLCAFPALSKLEFA-Ahx-Ahx-K(Aba)-$CONH_2$; peptide was synthesized in 5 µmol scale. After purification (column 1, 5-75% MeCN), the 3 x TFA salt product (5.2 mg, 1.91 µmol, 38% yield) was obtained as white solid. $t_R$ = 15.90 min. Purity ≥ 95%. Formula: $C_{112}H_{179}N_{27}O_{25}S_2$. Molecular weight: 2367.95. HRMS-ESI+ (m/z): $[M+3H]^{3+}$ calcd.: 790.1085; found: 790.1074.

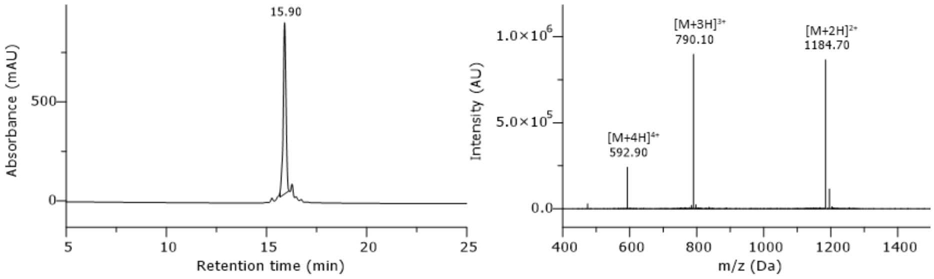

**Supplementary Fig. S5:** HPLC chromatogram of purified peptide $EPOP_{scr}$. Gradient 5-95% B in column 3, monitored at 220 nm.

**$EPOP_{wt}$-R8**

$H_2N$-EFSPLCLRALAFCALAK-Ahx-Ahx-K(Aba)-RRRRRRRR-$CONH_2$; peptide was synthesized in 20 µmol scale. After purification (column 2, 5-75% MeCN), the 11 x TFA salt product (18.1 mg, 3.72 µmol, 19% yield) was obtained as white solid. $t_R$ = 12.81 min. Purity ≥ 98%. Formula: $C_{160}H_{275}N_{59}O_{33}S_2$. Molecular weight: 3617.46. HRMS-ESI+ (m/z): $[M+4H]^{4+}$ calcd.: 905.2859; found: 905.2879.

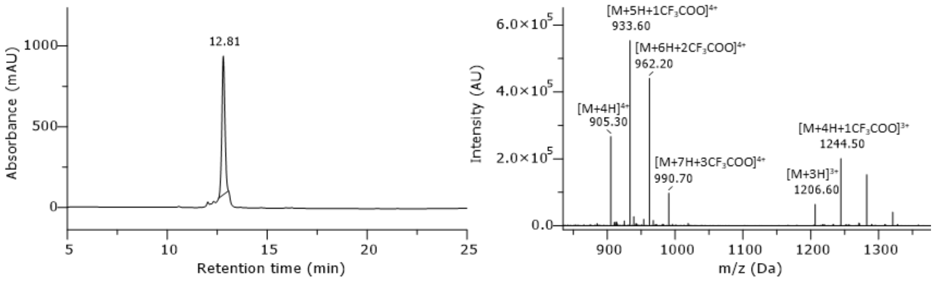

**Supplementary Fig. S6:** HPLC chromatogram of purified peptide $EPOP_{wt}$-R8. Gradient 5-95% B in column 3, monitored at 220 nm.

**$EPOP_{mut}$-R8**

$H_2N$-EFSPLC**A**RALAFCALAK-Ahx-Ahx-K(Aba)-RRRRRRRR-$CONH_2$; peptide was synthesized in 20 µmol scale. After purification (column 2, 5-75% MeCN), the 11 x TFA salt product (20.1 mg, 4.16 µmol, 21% yield) was obtained as white solid. $t_R$ = 12.20 min. Purity ≥ 97%. Formula: $C_{157}H_{269}N_{59}O_{33}S_2$. Molecular weight: 3575.38. HRMS-ESI+ (m/z): $[M+4H]^{4+}$ calcd.: 894.5277; found: 894.5236.

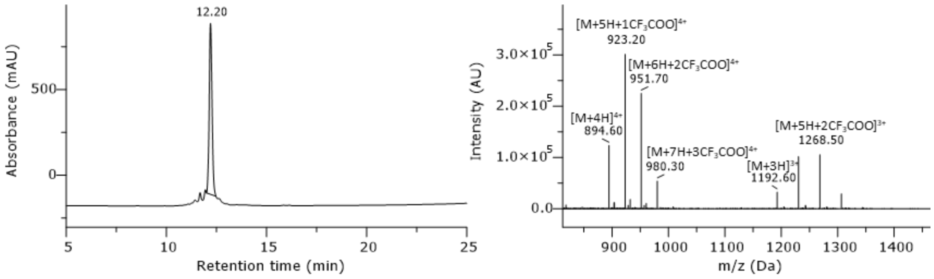

**Supplementary Fig. S7:** HPLC chromatogram of purified peptide $EPOP_{mut}$-R8. Gradient 5-95% B in column 3, monitored at 220 nm.

**EPOP$_{scr}$-R8**

H$_2$N-CLARLCAPFALSKLEFA-Ahx-Ahx-K(Aba)-RRRRRRRR-CONH$_2$; peptide was synthesized in 20 µmol scale. After purification (column 2, 5-75% MeCN), the 11 x TFA salt product (19.6 mg, 4.02 µmol, 20% yield) was obtained as white solid. $t_R$ = 11.58 min. Purity ≥ 99%. Formula: $C_{160}H_{275}N_{59}O_{33}S_2$. Molecular weight: 3617.46. HRMS-ESI+ (m/z): $[M+4H]^{4+}$ calcd.: 905.2859; found: 905.2898.

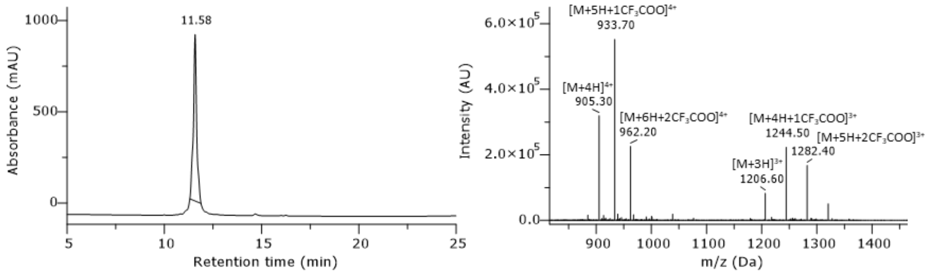

**Supplementary Fig. S8:** HPLC chromatogram of purified peptide EPOP$_{scr}$-R8. Gradient 5-95% B in column 3, monitored at 220 nm.

**R8**

H$_2$N-Ahx-Ahx-K(Aba)-RRRRRRRR-CONH$_2$; peptide was synthesized in 20 µmol scale. After purification (column 2, 5-75% MeCN), the 9 x TFA salt product (17.4 mg, 6.20 µmol, 31% yield) was obtained as white solid. $t_R$ = 6.95 min. Purity ≥ 96%. Formula: $C_{75}H_{140}N_{38}O_{13}$. Molecular weight: 1782.20. HRMS-ESI+ (m/z): $[M+2H]^{2+}$ calcd.: 891.5804; found: 891.5827.

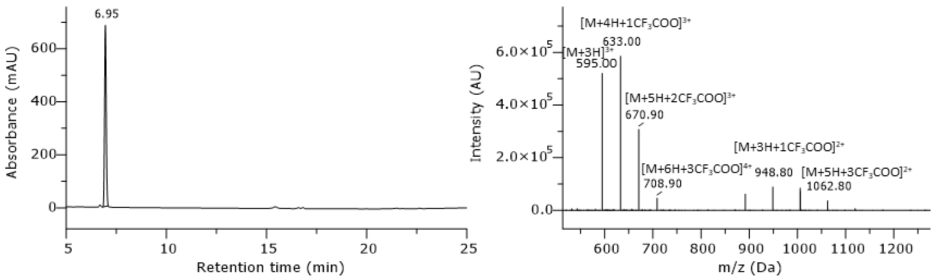

**Supplementary Fig. S9:** HPLC chromatogram of purified peptide R8. Gradient 5-95% B in column 3, monitored at 220 nm.

**Linker-R8$_{wt/mut}$ (resin-bound)**

Fmoc-K-Ahx-Ahx-K(Alloc)-RRRRRRRR-CONH$_2$; The resin-bound peptide was manually synthesized in 20 µmol scale. A test cleavage was performed to verify peptide identity before using it in automated SPPS. $t_R$ = 10.54 min. Purity ≥ 87%. Formula: $C_{91}H_{159}N_{39}O_{16}$. Molecular weight: 2055.53. HRMS-ESI+ (m/z): $[M+3H]^{3+}$ calcd.: 686.1024; found: 686.1033.

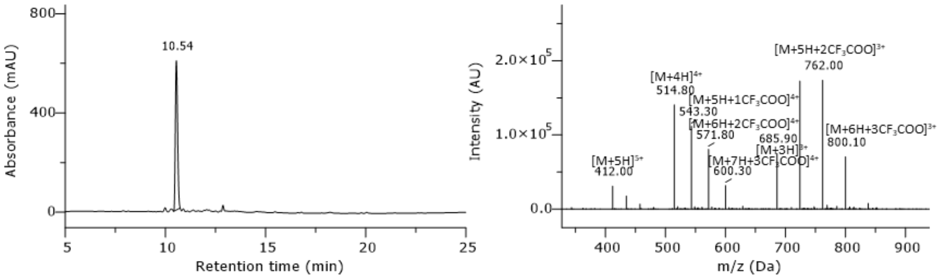

**Supplementary Fig. S10:** HPLC chromatogram of unpurified peptide R8-Resin$_{wt/mut}$. Gradient 5-95% B in column 3, monitored at 220 nm.

**Linker-R8$_{scr}$ (resin-bound)**

Fmoc-A-Ahx-Ahx-K(Alloc)-RRRRRRRR-CONH$_2$; The resin-bound peptide was manually synthesized in 20 µmol scale. A test cleavage was performed to verify peptide identity before using it in automated SPPS. $t_R$ = 11.65 min. Purity ≥ 90%. Formula: $C_{88}H_{152}N_{38}O_{16}$. Molecular weight: 1998.43. HRMS-ESI+ (m/z): $[M+3H]^{3+}$ cald.: 667.0831; found: 667.0839.

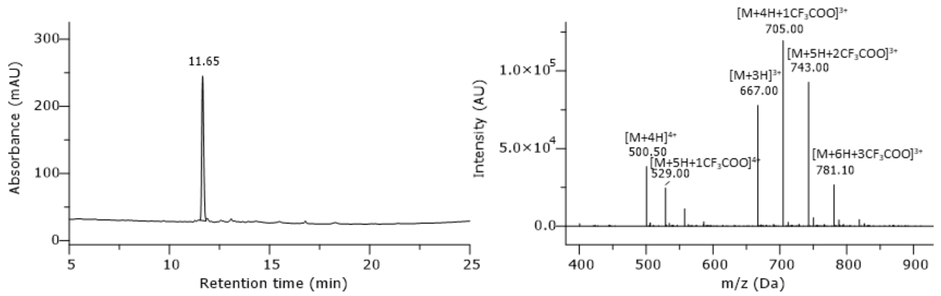

**Supplementary Fig. S11:** HPLC chromatogram of unpurified peptide R8-Resin$_{scr}$. Gradient 5-95% B in column 3, monitored at 220 nm.

#### 2.2 Hydrogen-Deuterium Exchange Mass Spectrometry (HDX-MS)

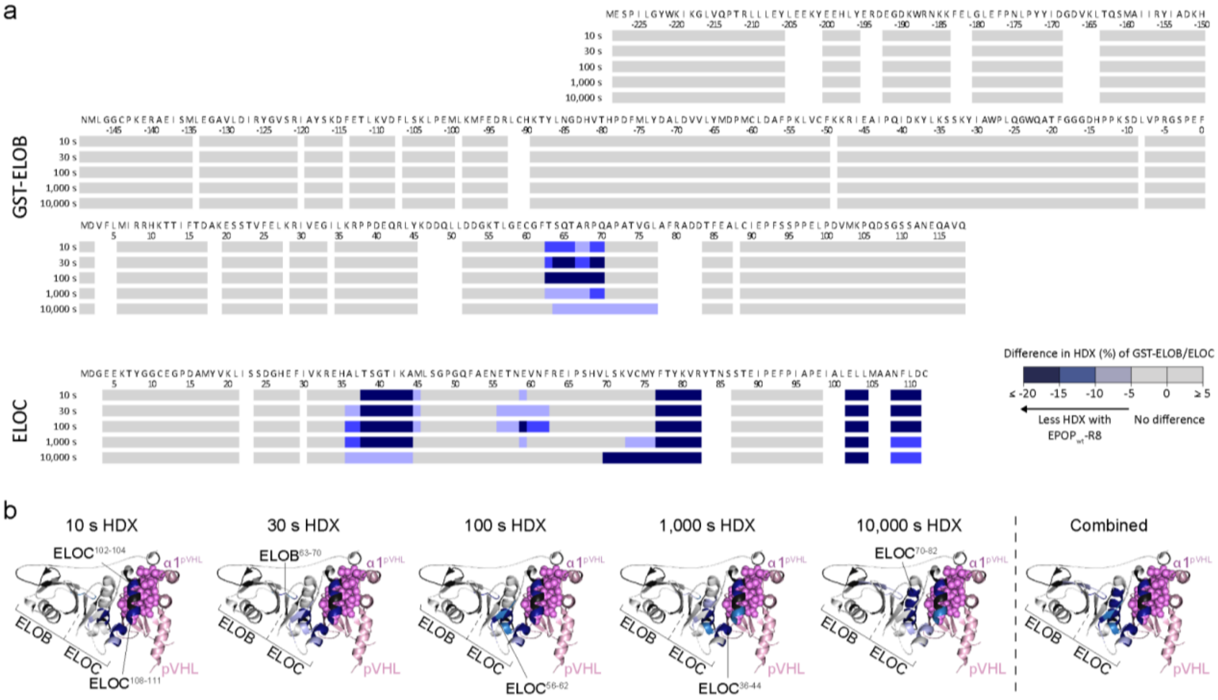

**Supplementary Fig. S12:** Conformational changes of GST-ELOB/ELOC induced by the EPOP$_{wt}$-R8 peptide. The difference in HDX of the GST-ELOB/ELOC complex in presence and absence of the EPOP$_{wt}$-R8 peptide is displayed (a) on the amino acid sequences of GST-ELOB (top) and ELOC (bottom), and (b) on the crystal structure of the HIF-1a/pVHL/ELOB/ELOC complex (PDB-ID 1LM8)[1]]. To render the residue-specific HDX differences from overlapping peptides in a for any given residue of GST-ELOB and ELOC, the shortest peptide covering this residue is employed. Where multiple peptides are of shortest length, the peptide with the residue closest to the peptide C-terminus is utilized. For residues not covered, no peptides could be obtained in HDX-MS (see source data). For the combined HDX differences in b, the HDX difference with the highest amplitude over all HDX timepoints is employed. Residues not covered by peptides are colored in black.

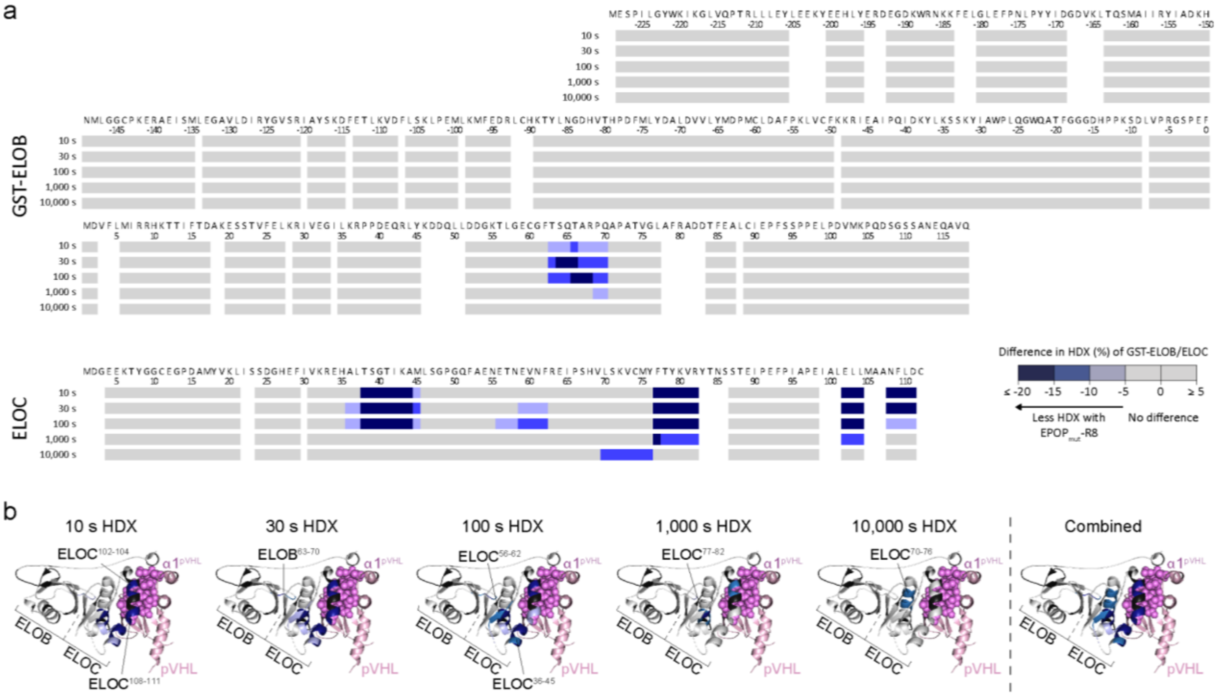

**Supplementary Fig. S13:** Conformational changes of GST-ELOB/ELOC induced by the EPOP$_{mut}$-R8 peptide. The difference in HDX of the GST-ELOB/ELOC complex in presence and absence of the EPOP$_{mut}$-R8 peptide is displayed (a) on the amino acid sequences of GST-ELOB (top) and ELOC (bottom), and (b) on the crystal structure of the HIF-1a/pVHL/ELOB/ELOC complex (PDB-ID 1LM8)[1]. To render the residue-specific HDX differences from overlapping peptides in a for any given residue of GST-ELOB and ELOC, the shortest peptide covering this residue is employed. Where multiple peptides are of shortest length, the peptide with the residue closest to the peptide C-terminus is utilized. For residues not covered, no peptides could be obtained in HDX-MS (see source data). For the combined HDX differences in b, the HDX difference with the highest amplitude over all HDX timepoints is employed. Residues not covered by peptides are colored in black.

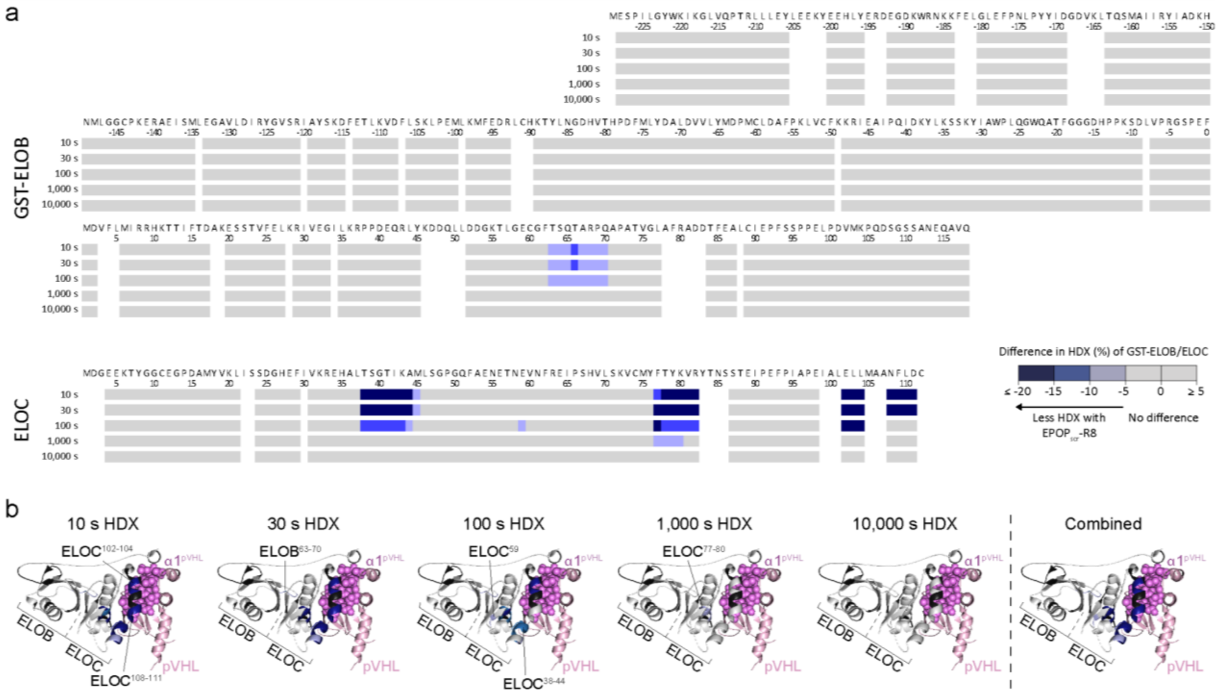

**Supplementary Fig. S14:** Conformational changes of GST-ELOB/ELOC induced by the EPOP$_{scr}$-R8 peptide. The difference in HDX of the GST-ELOB/ELOC complex in presence and absence of the EPOP$_{scr}$-R8 peptide is displayed (a) on the amino acid sequences of GST-ELOB (top) and ELOC (bottom), and (b) on the crystal structure of the HIF-1a/pVHL/ELOB/ELOC complex (PDB-ID 1LM8)[1]. To render the residue-specific HDX differences from overlapping peptides in a for any given residue of GST-ELOB and ELOC, the shortest peptide covering this residue is employed. Where multiple peptides are of shortest length, the peptide with the residue closest to the peptide C-terminus is utilized. For residues not covered, no peptides could be obtained in HDX-MS (see source data). For the combined HDX differences in b, the HDX difference with the highest amplitude over all HDX timepoints is employed. Residues not covered by peptides are colored in black.

#### 2.3 Cell Viability Assays

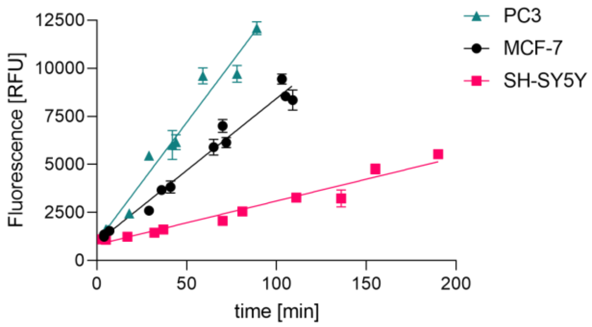

**Supplementary Fig. S15:** SH-SY5Y cells show lower signal intensities in resazurin based cell viability assays. For each cell line 20,000 cells have been seeded and read out as described for the resazurin-based cell viability assays. The fluorescence in relative fluorescence units (RFU) has been plotted over time. Data are represented as mean ± s.d. of n = 3 replicates.

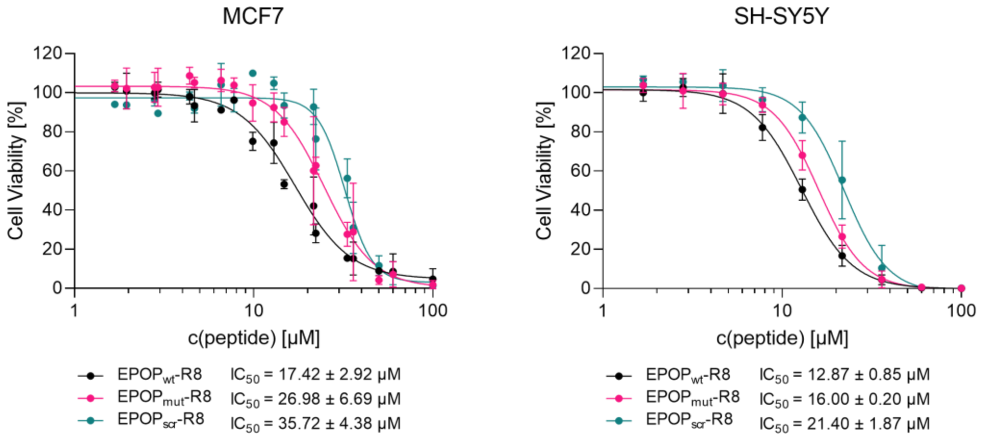

**Supplementary Fig. S16:** Dose-response curve of PC3, MCF7 and SH-SY5Y cells. All cell lines were treated with EPOP peptides for 24 h. Data from PC3 and MCF7 cells are based on resazurin assays. Data of SH-SY5Y cells are based on CellTiter-Glo®. Each plot represents mean ± s.d of n = 3 independent experiments.

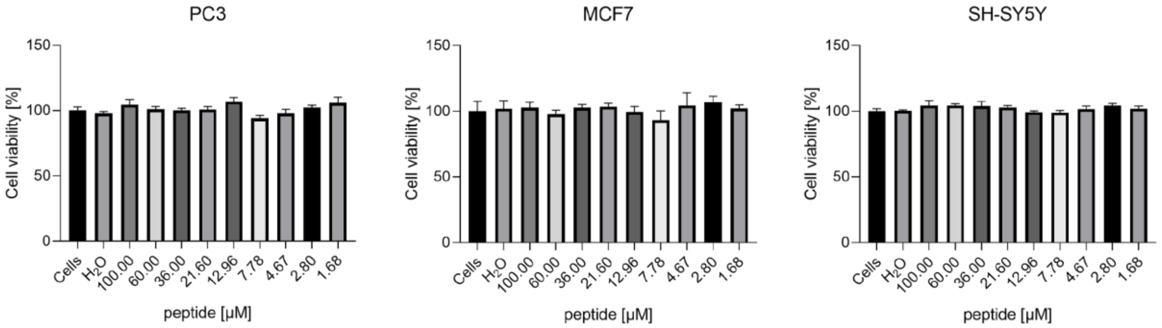

**Supplementary Fig. S17:** Dose-dependent toxicity of R8 peptide in PC3, MCF7 and SH-SY5Y cells after 24 h of incubation. All cell lines were treated with R8 peptides for 24 h. Data from PC3 and MCF7

cells are based on resazurin assays. Data of SH-SY5Y cells are based on CellTiter-Glo®. Each plot represents the mean ± s.d of n = 3 independent experiments.

#### 2.4 Replicates of apoptosis assays

**Replicate 1**

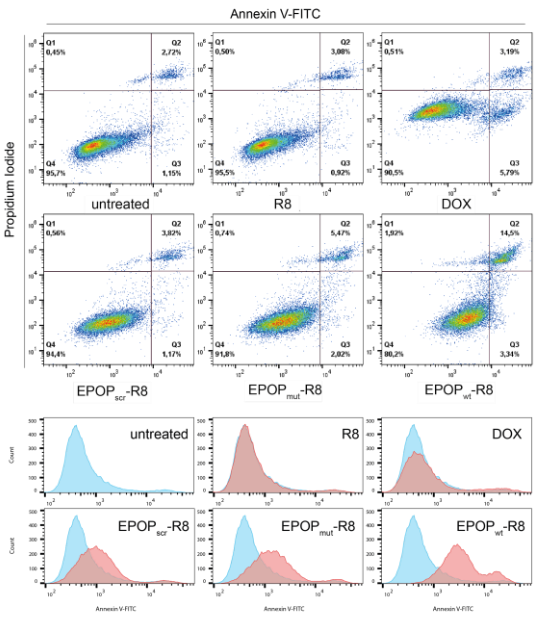

**Replicate 2**

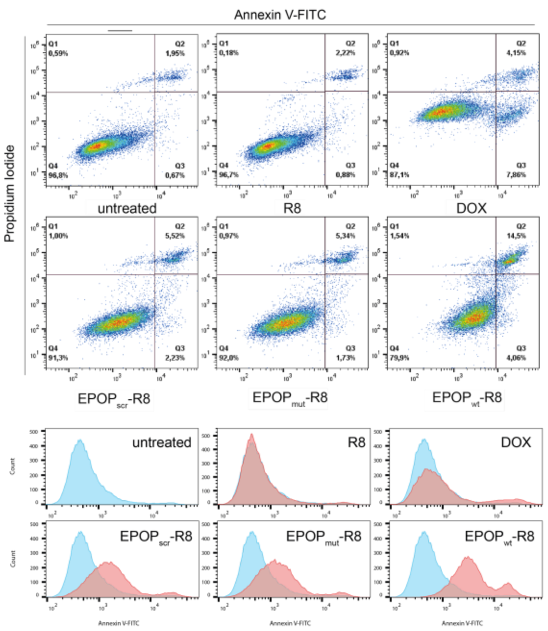

**Replicate 3**

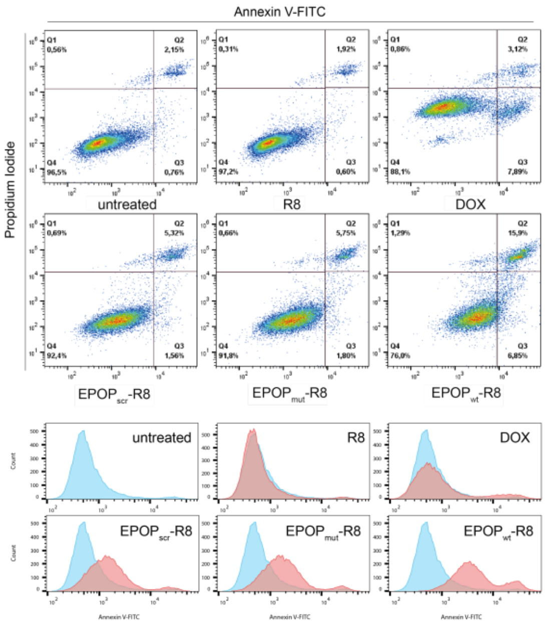

**Supplementary Fig. S18:** Flow cytometry experiments in PC3 cells. Results are derived from three independent experiments. The corresponding Annexin V-FITC histogram of each experiment has been plotted and shown below the dot plots.

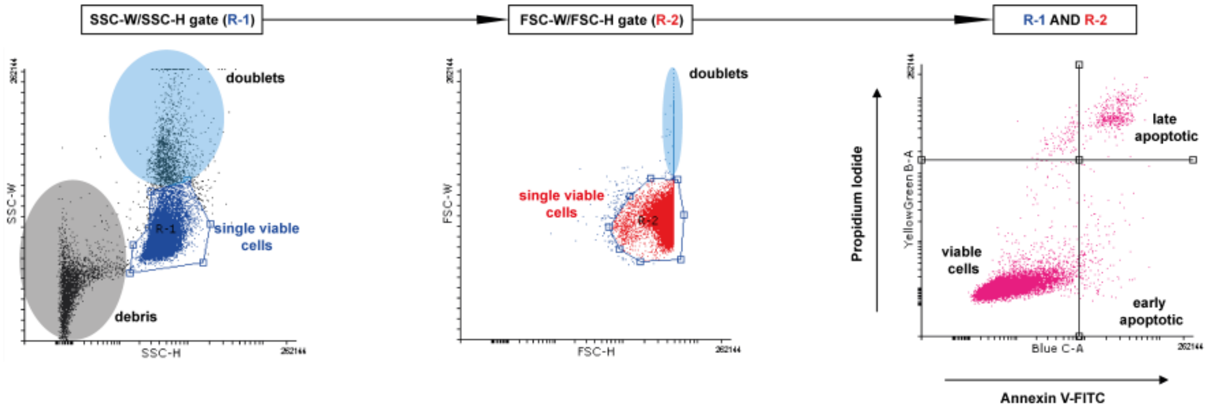

**Supplementary Fig. S19:** Hierarchical gating strategy for flow cytometry experiments with Annexin V-FITC and propidium iodide. Doublets and debris exclusion was performed by gating in a SSC-W vs SSC-H plot and a subsequent FSC-W vs FSC-H plot. The finale is shown as the propidium iodide (y-axis) vs Annexin V-FITC (x-axis).

#### 2.5 Silver for mass spectrometry

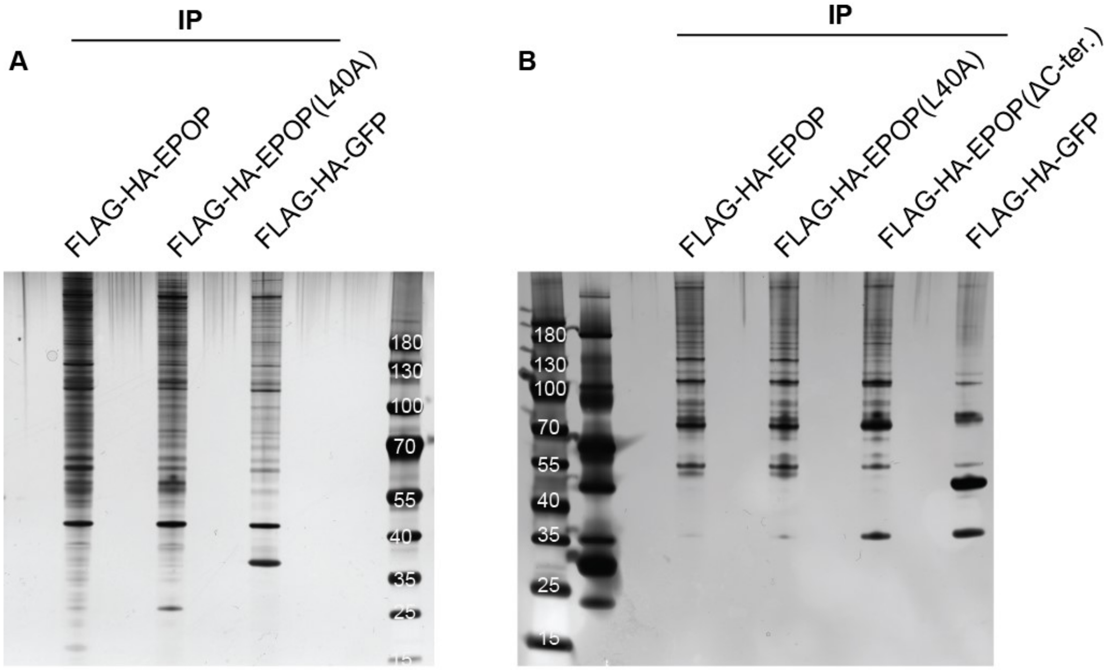

**Supplementary Fig. S20:** Silver stainings of co-immunoprecipitated material using various FLAG-EPOP proteins and FLAG-HA GFP (control) as bait. White numbers denote the molecular weight (in kDa) of the size standard.

#### 2.6 Replicates of Co-Immunoprecipitation experiments

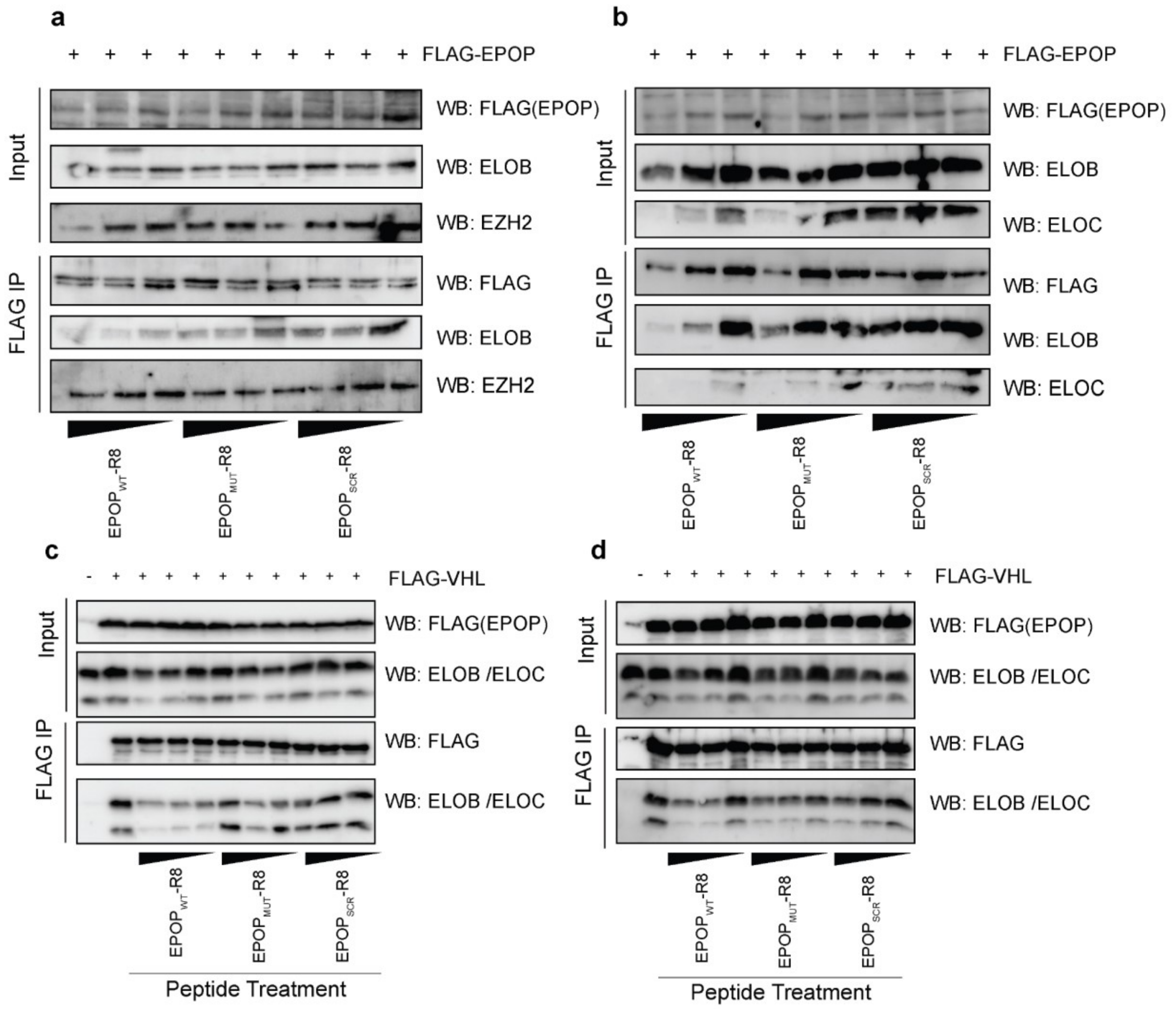

**Supplementary Fig. S21:** Additional replicates of co-immunoprecipitation experiments shown in Fig. 5c and d.

#### 2.7 Histograms of cell cycle analysis

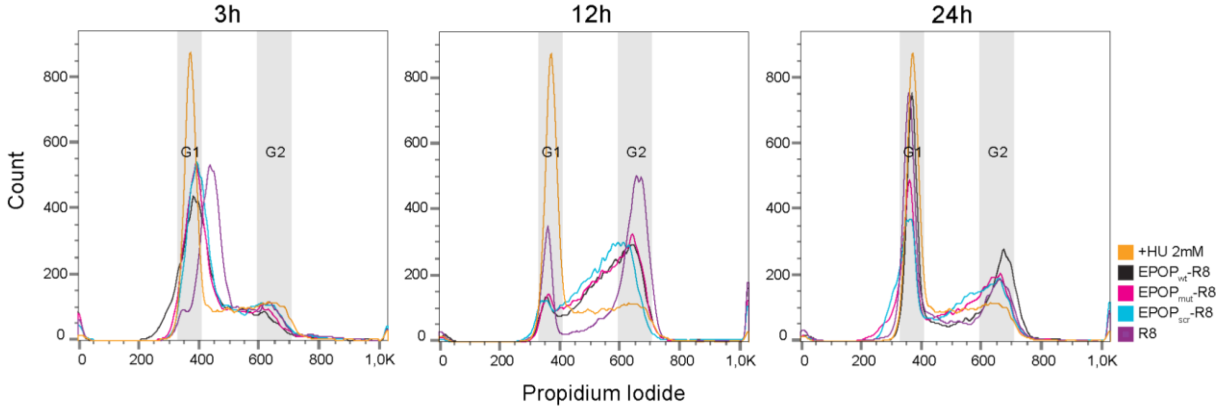

**Supplementary Fig. S22:** Representative histograms of cell cycle analysis shown in Ext. Data Fig. 5c.

#### 2.8 Antibodies and Oligonucleotides

**Supplementary Table S2:** Overview of antibodies used in this study.

| Antibodies | | |
|---|---|---|
| **Antigen** | **Source** | **Catalogue number** |
| **EloB[SF1] (rabbit)** | abcam | ab168836 |
| **EloC (rabbit)** | Bethyl | A304-787A |
| **EZH2** | Diagenode | C15410039 |
| **FLAG (mouse)** | Sigma Aldrich | F3165 |
| **SP-1 (rabbit)** | Self-made | Völkel et al. 2015 |
| **H3** | abcam | ab1791 |
| **HA (rat)** | Merck | 11867423 |
| **Suz12** | Active Motif | 103245 |
| **Beta Tubulin (KMX-1)** | Millipore | MAB3408 |
| **GST(B-14)** | Santa Cruz | sc-138 |
| **Mouse (sheep)HRP** | GE Healthcare | NA931 |
| **Rabbit (donkey)HRP** | GE Healthcare | NA934 |
| **Rabbit (goat) Alexa Fluor 488** | Thermo Fisher Scientific Inc. | A-11008 |
| **Rat (goat) HRP** | GE Healthcare | NA935 |
| **Rat(donkey) Alexa Fluor 488** | Thermo Fisher Scientific Inc. | A-21208 |
| **Rat (goat) Alexa Fluor 546** | Thermo Fisher Scientific Inc. | A-11081 |

**Supplementary Table S3:** Overview of oligonucleotides used in this study.

| Oligonucleotides | sequence | Source |
|---|---|---|
| **shRNA Control sequence** | CAACAAGATGAAGAGCACCAA | SHC002 (Sigma Aldrich) |
| **shRNA targeting sequence ELOB:** | GTGTGGCTTCACCAGTCAAAC | Liefke et al. 2016 |
| **shRNA targeting sequence ELOC** | GAAACCAATGAGGTCAATTT | Liefke et al. 2016 |
| **RT hGAPDH for** | AGCCACATCGCTCAGACAC | This paper |

| | | |
|---|---|---|
| **RT hGAPDH rev** | GCCCAATACGACCAAATCC | This paper |
| **RT hELOC for** | CATCAGGCACGATAAAAGCCA | This paper |
| **RT hELOC rev** | GCTGTTAGTGTAGCGAACCTTG | This paper |
| **RT hELOB for** | CGAACTGAAGCGCATCGTC | This paper |
| **RT hELOB rev** | TCCAAGAGTTGGTCATCCTTGT | This paper |

1. Min, J.H. et al. Structure of an HIF-1alpha -pVHL complex: hydroxyproline recognition in signaling. *Science* **296**, 1886-9 (2002).
